# Fitness effects of CRISPR endonucleases in *Drosophila melanogaster* populations

**DOI:** 10.1101/2021.05.13.444039

**Authors:** Anna M. Langmüller, Jackson Champer, Sandra Lapinska, Lin Xie, Matthew Metzloff, Jingxian Liu, Yineng Xu, Andrew G. Clark, Philipp W. Messer

## Abstract

CRISPR/Cas9 systems provide a highly efficient and flexible genome editing technology with numerous potential applications in areas ranging from gene therapy to population control. Some proposed applications involve CRISPR/Cas9 endonucleases integrated into an organism’s genome, which raises questions about potentially harmful effects to the transgenic individuals. One application where this is particularly relevant are CRISPR-based gene drives, which promise a mechanism for rapid genetic alteration of entire populations. The performance of such drives can strongly depend on fitness costs experienced by drive carriers, yet relatively little is known about the magnitude and causes of these costs. Here, we assess the fitness effects of genomic CRISPR/Cas9 expression in *Drosophila melanogaster* cage populations by tracking allele frequencies of four different transgenic constructs, designed to disentangle direct fitness costs due to the integration, expression, and target-site activity of Cas9 from costs due to potential off-target cleavage. Using a maximum likelihood framework, we find a moderate level of fitness costs due to off-target effects but do not detect significant direct costs. Costs of off-target effects are minimized for a construct with Cas9HF1, a high-fidelity version of Cas9. We further demonstrate that using Cas9HF1 instead of standard Cas9 in a homing drive achieves similar drive conversion efficiency. Our results suggest that gene drives should be designed with high-fidelity endonucleases and may have implications for other applications that involve genomic integration of CRISPR endonucleases.

## Introduction

The ability to make specific edits of genetic material has been a long-standing goal in molecular biology. Until recently, such DNA engineering was cumbersome, expensive, and difficult since it relied on site-specific nucleases or random insertions. CRISPR technology represents a milestone in genome editing because it makes DNA engineering highly efficient, relatively simple to use, and cost-effective through the use of endonucleases that can be flexibly programmed to cut specific sequences dictated by a guide RNA (gRNA) (1, 2).

The programmability of CRISPR/Cas9 systems allows for numerous potential applications (1), including cancer and disease treatment (3–7), stimuli tracking in living cells (8), and crop improvement (9). While most applications of CRISPR use this technology to engineer specific modifications in a given gene sequence, some proposed applications take the idea one step further by integrating the CRISPR machinery itself into an organism’s genome. In that case, endonuclease activity can continue to produce genetic changes in the cells of the living organism. When present in the germline, these genetic changes might even be passed on to future generations, such as in CRISPR-based gene drives — “selfish” genetic elements that are engineered to rapidly spread a desired genetic trait through a population into which they are released (10–14).

However, major questions loom large about the technical feasibility of these proposed applications. For example, it remains unclear whether activity of CRISPR endonucleases could entail unintended and potentially harmful consequences in the transgenic organisms, for instance due to the tendency to produce non-specific DNA modifications (so-called “off-target effects”) (15). Such off-target cleavage could be substantially higher when Cas9 is continuously expressed from a genome and inherited by offspring, where further off-target cleavage can occur.

In this study, we seek to address this question in the context of CRISPR gene drive, a new technology that could potentially be used for applications ranging from the control of vector-borne diseases to the suppression of invasive species (10, 12, 14, 16). One class of CRISPR-based gene drives are so-called “homing drives”. These genetic constructs are programmed to cleave a wild-type sister chromatid and get copied to the target site through homology-directed repair. Since “homing” occurs in the germline, the drive allele will be inherited at a super Mendelian rate and can thereby spread quickly through the population. The effectiveness of such systems has now been demonstrated in various organisms, including yeast (17–20), mosquitoes (21–24), fruit flies (25–32), and mice (33). Another class of CRISPR gene drives operate by the “toxin-antidote” principle (34). Here, the drive allele serves as the “toxin” by carrying a CRISPR endonuclease programmed to target and disrupt an essential wild-type gene. At the same time, the construct also contains a recoded version of that gene (the “antidote”), which is immune to cleavage by the drive. Over time, such a drive will continuously remove wild-type alleles from the population, while the drive allele will increase in frequency (35). Both homing and toxin-antidote drives can be “modification drives”, intended to spread a desired genetic payload through the population (e.g., a gene that prevents mosquitoes from transmitting malaria), or “suppression drives”, where the goal is to diminish or outright eliminate the target population (34, 36).

A key factor in determining the expected population dynamics of any type of gene drive is the fitness cost imposed by the drive (37). Such fitness costs could come in the form of reduced viability, fecundity, or mating success of the individuals that carry drive alleles. In suppression drives, some fitness costs are typically an intended feature of the drive, necessary to ultimately achieve population suppression. However, these costs are usually recessive to allow the drive to spread to high frequency, and there is generally a limit as to how high other costs can be before the drive will lose its ability to spread effectively (34, 36, 38, 39). For modification drives, fitness costs tend to slow the spread of the drive and can thereby increase the chance that resistance alleles evolve, which could ultimately outcompete the drive (12). For such applications, it is therefore desirable to minimize any fitness costs. In drives with frequency-dependent invasion dynamics, such as most CRISPR toxin-antidote systems (32, 40), fitness costs can further determine the frequency threshold required for the drive to spread through the population (34, 36, 38).

We believe it is useful to distinguish between two types of fitness costs of a gene drive. The first class of “direct” costs comprise any effects resulting from the genomic integration of the drive construct (e.g., when this disrupts a functionally important region), costs of potential “payload” genes included in the drive construct, costs resulting directly from the expression of the endonuclease or other drive elements such as gRNAs, and costs due to cleavage of the intended target site. The second class comprise any potential fitness costs due to “off-target” activity of the CRISPR endonuclease, referring to cleavage and disruption of any unintended sites in the genome. Despite their critical importance, we still know surprisingly little about the specific types of fitness costs imposed by gene drives. In particular, it remains unclear whether there are certain “baseline” fitness costs that would be difficult to avoid in any gene drive construct, for instance because they are inherent to the expression and activity of the CRISPR endonuclease or result from their tendency to generate off-target effects.

In this study, we conduct a comprehensive assessment of the fitness effects resulting from genomic expression of CRISPR/Cas9 in experimental *Drosophila melanogaster* populations. We specifically investigate four different transgenic constructs that allow us to disentangle direct fitness costs from those due to off-target effects. We estimate these fitness costs both through a statistical analysis of allele frequency trajectories in cage populations and a direct evaluation of individual fitness components using viability, fecundity, and mate choice assays.

## Results

### Construct design

We designed four constructs to assess the fitness costs of *in vivo* CRISPR/Cas9 expression in *D. melanogaster*. As a starting point for our transgenic fly lines, we engineered an EGFP fluorescent marker driven by the 3xP3 promoter into a gene-free, non-heterochromatic position on chromosome 2L (region targeted by gRNA: 20,368,542 - 20,368,561; Figure 1A). This EGFP marker was then used as insertion point for the four constructs we tested. Our first construct, “Cas9_gRNAs”, contains Cas9 expressed by the *nanos* promoter, the fluorescence marker DsRed driven by the 3xP3 promoter, and four gRNAs driven by the U6:3 promoter (Figure 1B), which are separated by tRNAs that are removed after transcription (29). The gRNAs of the Cas9_gRNAs construct target a gene-free, non-hetero-chromatic position on a different chromosome (3L, region targeted by gRNAs: 18,297,270 – 18,297,466), preventing any homing activity. In addition to Cas9_gRNAs, three other constructs were designed: “Cas9_no-gRNAs” has a similar architecture as Cas9_gRNAs, but lacks the four gRNAs driven by the U6:3 promoter (Figure 1C); “no-Cas9_no-gRNAs” contains neither Cas9, nor the gRNAs, but only the fluorescence marker DsRed driven by the 3xP3 promoter (Figure 1D); the last construct, “Cas9HF1_gRNAs” (Figure 1E), has the same architecture as Cas9_gRNAs, except that Cas9 is replaced by a high-fidelity version (Cas9HF1), which has been reported to largely eliminate off-target cleavage (41). As expected, all progeny of individuals with the Cas9_gRNAs and Cas9HF1_gRNAs alleles had at least one of their gRNA target sites mutated, together indicating that all four gRNAs were active in both these constructs.

**Figure 1.**
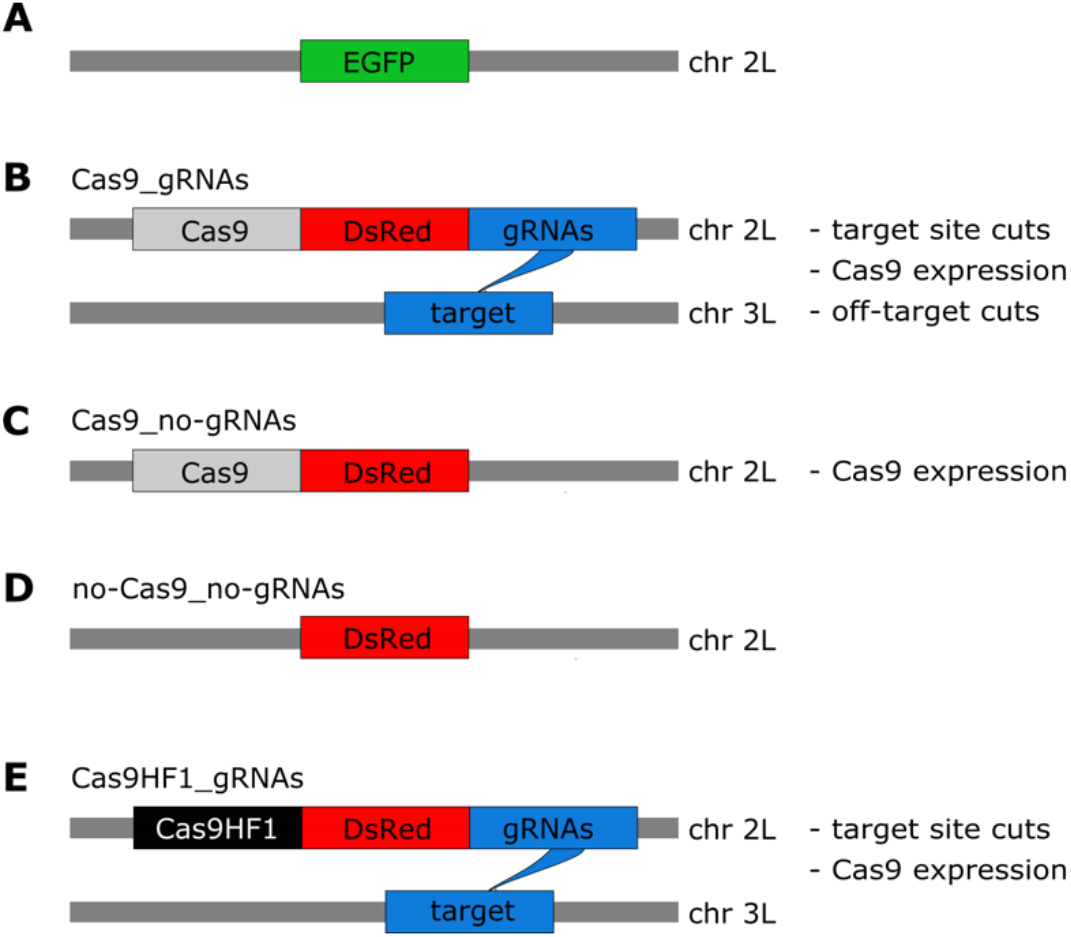
Overview of constructs and the potential types of fitness costs in the four constructs. (A) The starting point for our constructs is an EGFP marker inserted into chromosome 2L (~20.4 Mb). The four constructs are then inserted into this EGFP locus (thereby disrupting EGFP). (B) The Cas9_gRNAs construct contains Cas9, DsRed, and gRNAs. The gRNAs target chromosome 3L (~18.3 Mb), instead of the sister chromatid. (C) The Cas9_no-gRNAs construct carries Cas9 and DsRed, but no gRNAs are expressed. (D) The no-Cas9_no-gRNAs construct carries only the fluorescent marker DsRed. (E) The Cas9HF1_gRNAs construct has the same structure as Cas9_gRNAs but carries Cas9HF1 instead of Cas9.

The specific designs of these four different constructs allow us to identify and disentangle different types of Cas9-related fitness costs. If double strand breaks at the target site impose fitness costs, such costs should be present for the Cas9_gRNAs and Cas9HF1_gRNAs constructs, but not for the Cas9_no-gRNAs and no-Cas9_no-gRNAs constructs, since Cas9_no-gRNAs has no gRNAs expressed to guide Cas9 to the target site, and the no-Cas9_no-gRNAs construct neither expresses Cas9 nor the gRNAs. If the expression of Cas9 imposes a fitness cost, all constructs except for no-Cas9_no-gRNAs should incur such a cost, because only this construct does not express Cas9. If off-target effects of Cas9 impose fitness costs, only the Cas9_gRNAs construct should incur them, because the designs of Cas9_no-gRNAs and no-Cas9_no-gRNAs prevent cutting events, and Cas9HF1_gRNAs reportedly has a much lower rate of off-target cleavage (41). Figure 1 summarizes the designs and different potential fitness costs for our four constructs.

### Population cage experiments

To assess the fitness effects of the four constructs, we tracked their population frequencies relative to the baseline EGFP construct over several generations in large cage populations. Overall, we assessed 13 cages: seven with the Cas9_gRNAs construct, and two each with the Cas9_no-gRNAs, no-Cas9_no-gRNAs, and Cas9HF1_gRNAs construct (Figure 2). In each cage population, the construct frequency was tracked for at least eight consecutive, non-overlapping generations. The median population size across all experiments was 3,602 (Figure S1 & Supplementary Results). To avoid potentially confounding maternal fitness effects on the construct frequency dynamics, we excluded the first generation of five cage populations (Cas9_gRNAs construct: replicates 1, 2, 5, 6, and 7) from the analysis, because their founding construct homozygotes and EGFP homozygotes were raised in potentially different environments.

**Figure 2.**
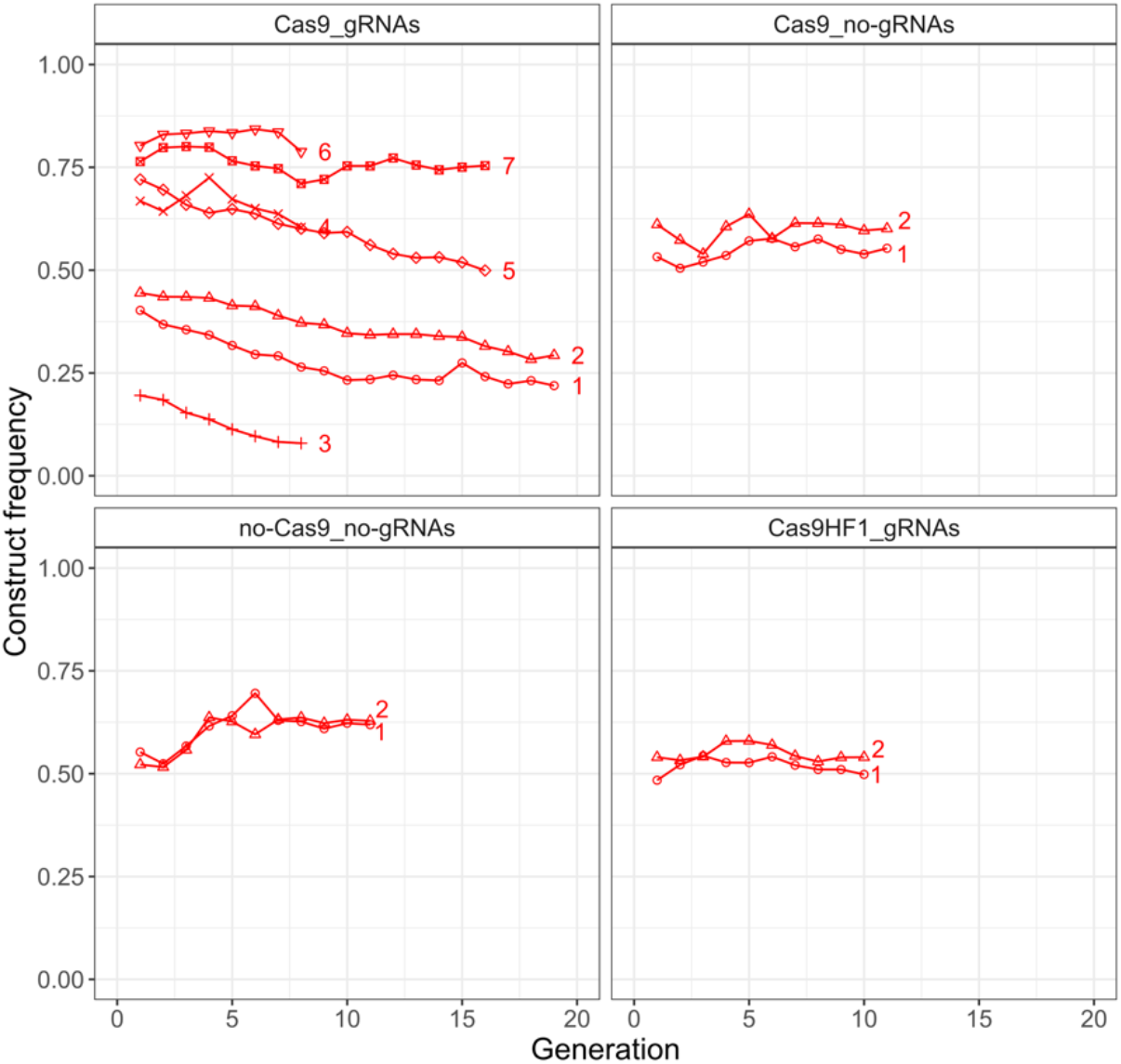
Construct frequency trajectories in the cage populations. Each line is one cage experiment.

We found Cas9_gRNAs to be the only construct that systematically decreased in frequency across all replicate cages (Figure 2). Interestingly, the allele frequency change was not consistent with fixed direct fitness costs. Instead, the construct frequency “bottomed out” in most replicates, and this occurred more quickly when the starting frequency was higher (Figure 2). We do not expect that the different frequency trajectories were caused by replicate-specific maternal effects, since replicates 3 and 4, which had very different frequency dynamics and starting frequencies, originated from the same pool of founding construct and EGFP homozygotes. In contrast to Cas9_gRNAs, the three other constructs did not decrease in frequency consistently across replicates, suggesting Cas9 off-targets effects are the primary driver of the fitness costs we detected (Figure 1).

A model in which fitness costs are predominantly caused by a limited set of potential off-target sites (e.g., because they have similar sequence to the actual target site) also suggests a possible mechanism for the observed “bottoming out” of the Cas9_gRNA frequency trajectories. At the beginning of the experiments, all genomes in individuals without the CRISPR construct will harbor uncut alleles at all these sites. In individuals carrying the construct, such sites may be cut and then repaired by end joining, which typically results in a mutated sequence. Some of these mutations could be deleterious (e.g., if they change the sequence of an important gene). A mutated site will also be protected to future cutting, similar to the creation of resistance alleles in a homing drive (28). Early in the experiment, mutated off-target sites will be found primarily in individuals that also carry drive alleles. This will lower the fitness of these individuals and, consequently, impose negative selection against construct alleles. However, as mutated off-target sites accumulate over the course of an experiment, they will increasingly segregate independently from construct alleles, thereby reducing selection against these alleles. By the time all potential off-target sites in the population have been cut, construct alleles would no longer experience any negative selection if such off-target effects were indeed the only mechanism underlying the construct’s fitness costs. Due to the higher overall rate of cleavage events in the population, cages where the construct is introduced at a higher frequency will experience this effect faster than cages where it is introduced at lower frequency, consistent with our experiments.

### Maximum likelihood analysis

To quantify the fitness costs of the different constructs from the observed frequency trajectories in our experiments, we adopted a previously developed maximum likelihood framework (42) and modified it to model two unlinked autosomal loci, representing the construct and a single idealized off-target site (see Methods). Each of the two loci is biallelic (EGFP/construct; uncut/cut off-target site). In individuals that carry a construct, all uncut off-target alleles are assumed to be cut in the germline, which are then passed on to offspring that could suffer negative fitness consequences. In the early embryo, all uncut off-target alleles are assumed to be cut by maternally deposited Cas9 if the mother carries at least one construct allele, changing the individual’s genotype at the off-target site and exposing it to the potential fitness costs associated with this new genotype. Fitness costs due to carrying the construct and/or the presence of cut off-target sites are assumed to be multiplicative across the two loci, as well as for the two alleles at each locus. We studied models where fitness costs affect only viability, and models where they affect only mate choice and fecundity (both equally). Overall, our maximum likelihood model infers three parameters: the effective population size *N_e_*, the “direct fitness estimate” (defined as the relative fitness of construct/EGFP heterozygotes versus EGFP/EGFP homozygotes), and the “off-target fitness estimate” (defined as the relative fitness of cut/uncut heterozygotes versus uncut/uncut homozygotes.). Note that in our idealized model with a single cleavage site, this site could in principle also represent “on-target” cleavage. However, due to the intergenic location of all gRNA target sites in our constructs, we do not expect such fitness costs to be present. Furthermore, if on-target cleavage had a measurable negative fitness effect, this should have been apparent in the frequency trajectories of the Cas9HF1_gRNAs construct. Since this construct had no apparent reduction in fitness, we refer to this fitness parameter as exclusively “off-target”.

For each construct, five different models were studied: In the “full inference model”, both the construct and cut off-target alleles can impose fitness costs. In the “construct” model, only construct alleles impose a fitness cost. In the “off-target” model, only cut off-target alleles impose a fitness cost. In the “initial off-target model”, we assumed that fitness costs originated before the experiment (e.g., through the injection process or maternal effects in the ancestral generation). For the “initial off-target model”, the construct homozygotes in the ancestral generation all had cut off-target alleles, but no additional off-target cutting occurred during the experiment (i.e., the germline and embryo cut rate were set to 0). Finally, in the “neutral” model, no fitness costs were present at all. Inferences were performed on the combined data of the replicated experimental populations for each construct. The individual models were compared using the corrected Akaike Information Criterion (AICc) (43) — a goodness-of-fit measure that also penalizes for complexity (i.e., number of parameters) in a given model. A lower AICc value indicates a higher quality model.

Table 1 shows the results for the Cas9_gRNAs construct. Here, we found the full inference model with viability selection to yield the highest quality, with a “direct fitness estimate” of 0.98 and an “off-target fitness estimate” of 0.84. Note, however, that the 95% confidence interval of the direct fitness estimate includes 1, and the simpler “off-target” model with viability selection in which the direct fitness estimate is set to 1 in fact has an equal AICc value to the “full” model. Thus, direct fitness costs are likely smaller than 5% in construct/EGFP heterozygotes. Models with fecundity/mate choice selection generally had lower quality than models with viability selection. The “initial off-target” and “neutral” models yielded the lowest AICc values. Taken together, these results suggest that the observed frequency trajectories of the Cas9_gRNAs construct in our cage populations are best explained by a model where direct effects are less than a few percent and off-target effects impose moderate fitness costs of ~30% (= 1-0.84^2^) in cut/cut homozygotes in our idealized single off-target site model.

**Table 1.** Model comparison and parameter estimates for Cas9_gRNAs.

| model | selection | $\hat{N}_e$ | direct fitness<br>estimate | off-target fitness<br>estimate | $\ln\hat{L}$ | $P$ | AICc |
| --- | --- | --- | --- | --- | --- | --- | --- |
| full | viability | 175<br>[140 – 215] | 0.98<br>[0.95 – 1.00] | 0.84<br>[0.77 – 0.91] | 384.7 | 3 | -763 |
| full | mate choice<br>= fecundity | 163<br>[131 – 200] | 0.96<br>[0.94 – 0.98] | 1.00<br>[0.95 – 1.06] | 378.8 | 3 | -751 |
| construct | viability | 164<br>[131 – 201] | 0.96<br>[0.93 – 0.98] | 1* | 378.9 | 2 | -754 |
| construct | mate choice=<br>fecundity | 163<br>[131 - 200] | 0.96<br>[0.94 – 0.98] | 1* | 378.8 | 2 | -754 |
| off-target | viability | 173<br>[139 - 212] | 1* | 0.80<br>[0.74 – 0.88] | 383.6 | 2 | -763 |
| off-target | mate choice<br>= fecundity | 157<br>[126 - 192] | 1* | 0.95<br>[0.90 – 1.01] | 375.1 | 2 | -746 |
| initial off-target | viability | 156<br>[125 - 191] | 1* | 0.92<br>[0.82 – 1.02] | 374.8 | 2 | -745 |
| initial off-target | mate choice<br>= fecundity | 156<br>[125 - 191] | 1* | 0.96<br>[0.91 – 1.01] | 374.8 | 2 | -745 |
| neutral | none | 154<br>[123 – 189] | 1* | 1* | 373.6 | 1 | -745 |

Each row shows the parameter estimates ( $\hat{N}_e$ = effective population size), maximum log Likelihood ( $\ln\hat{L}$ ), number of free parameters in the maximum likelihood framework ( $P$ ), and corrected Akaike Information Criterion value ( $AICc = 2p - 2\ln\hat{L} + (2p^2 + 2p)/(n - p - 1)$ where $n = 87$ is the number of generation transitions) for a specific model and selection type. 1\* entries indicate that a parameter was fixed at 1 (= no fitness effect is estimated). Values in squared brackets in the parameter estimate columns represent the 95 % confidence intervals estimated from a likelihood ratio test with one degree of freedom.

To test whether this model can accurately capture the observed frequency-dependent construct dynamics of the Cas9_gRNAs construct, we simulated construct trajectories under the model with the best AICc value (full inference model, viability selection) under its maximum likelihood parameter estimates (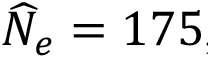, direct fitness estimate = 0.98, off-target fitness estimate = 0.84). The simulations do not only resemble the observed decrease in construct frequency, but also capture the bottoming out of individual replicates depending on their construct starting frequency (Figure 3). Additionally, we compared simulated trajectories for this model with simulated trajectories from the “construct” model with viability selection (Figure S2). We found that the full inference model captures the observed frequency dependent construct dynamics better than the model that only considers direct fitness costs, with most of the improvement due to better matching trajectories from cages with low starting frequencies, where off-target effects would be expected to have a more drastic impact on the relative fitness of construct-carrying individuals.

**Figure 3.**
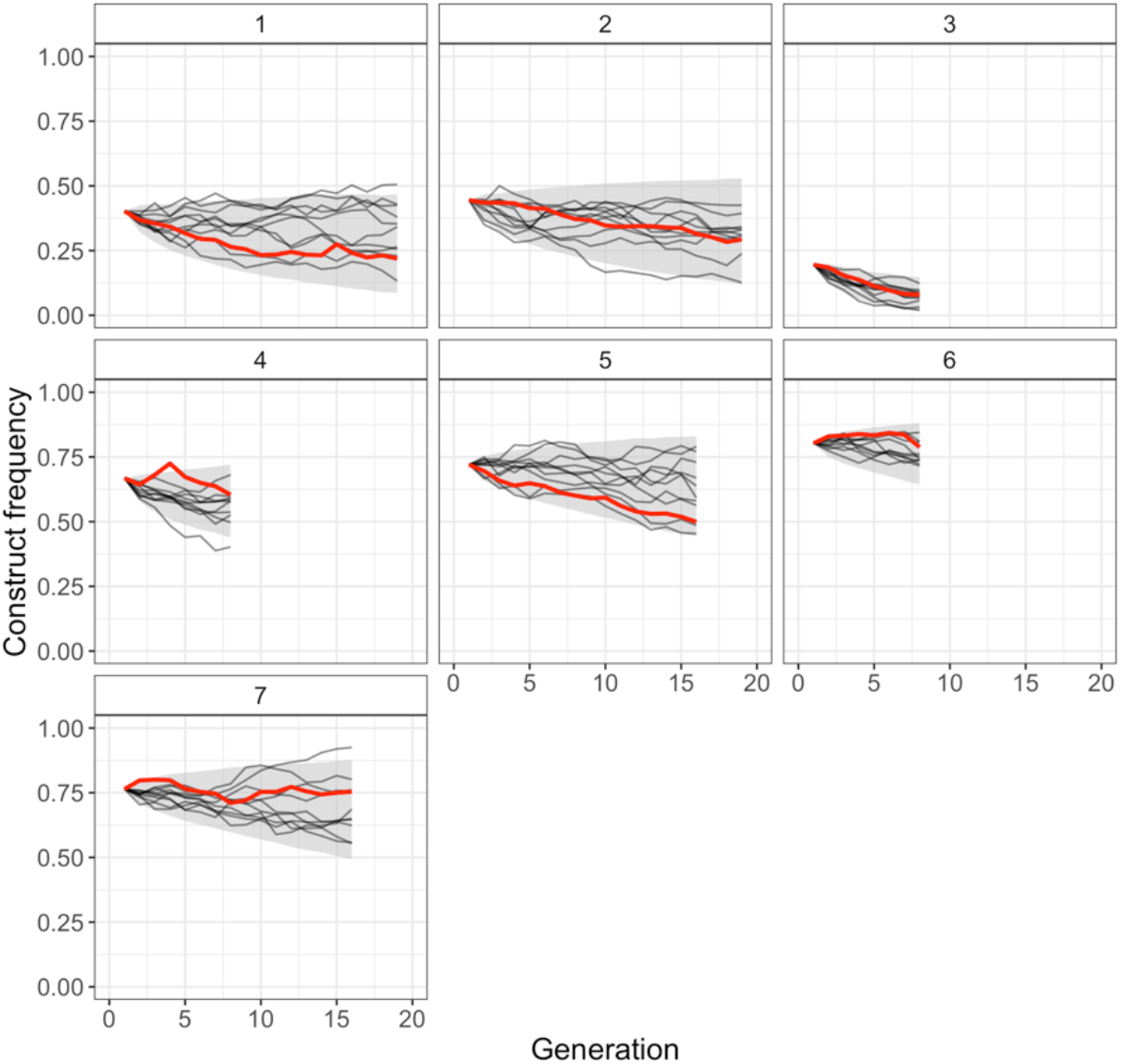
Comparison of observed Cas9_gRNAs construct frequencies with simulated trajectories of a full model with viability selection (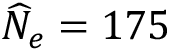, direct fitness estimate = 0.98, off-target fitness estimate = 0.84). Solid red lines present observed construct frequencies, black lines show ten simulated trajectories for each cage, and the shaded area represents the range between the 2.5 and 97.5 percentile of the simulated trajectories (10,000 simulations per cage).

To further support our hypothesis that fitness costs are primarily driven by off-target effects, we applied the maximum likelihood inference framework to the experimental cage data of the three other constructs (Cas9_no-gRNAs, no-Cas9_no-gRNAs, and Cas9HF1_gRNAs). Because none of these three constructs should be capable of producing substantial amounts of off-target cuts by design, we set the germline and embryo cut rate to 0 and inferred viability fitness effects for the construct. Except for the “initial off-target” model, construct homozygotes of the ancestral population were assumed to not carry any cut off-target alleles. For Cas9_no-gRNAs, and no-Cas9_no-gRNAs, the “neutral” model without any fitness costs explains the observed construct frequency trajectories best (Table 2, Figure S3), corroborating the notion that off-target cuts are the main driver of Cas9 fitness costs in our experimental populations (Figure 1). However, the construct frequency dynamics of Cas9HF1_gRNAs are best explained by an “initial off-target” model, where cut off-target alleles are beneficial, closely followed by the neutral model (Table 2). While we cannot rule out that the initial construct homozygotes of Cas9HF1_gRNAs had a fitness advantage due to cut off-target alleles or transgenerational beneficial effects, the 95% confidence interval for the off-target fitness parameter is broad and includes 1. This putative fitness advantage could also potentially be explained by maternal effects that persisted for 2-3 generations. Although we do not anticipate that any other construct than Cas9_gRNAs can produce substantial off-target effects, we repeated the analysis of the three other constructs with cut rates set to 1 and inferred viability selection, which yielded similar results (Table S1).

**Table 2.** Model comparison and parameter estimates for Cas9_no-gRNAs, no-Cas9_no-gRNAs, and Cas9HF1_gRNAs.

| construct | model | selection | $\hat{N}_e$ | direct fitness<br>estimate | off-target<br>fitness estimate | $\ln\hat{L}$ | $P$ | AICc |
| --- | --- | --- | --- | --- | --- | --- | --- | --- |
| Cas9_no-gRNAs | construct | viability | 243<br>[152 – 366] | 1.0<br>[0.96 – 1.04] | 1* | 88.6 | 2 | -173 |
| Cas9_no-gRNAs | initial<br>off-target | viability | 250<br>[156 - 377] | 1* | 0.84<br>[0.65 – 1.18] | 89.2 | 2 | -174 |
| Cas9_no-gRNAs | neutral | none | 243<br>[152 – 366] | 1* | 1* | 88.6 | 1 | -175 |
| no-Cas9_no-gRNAs | construct | viability | 162<br>[101 – 243] | 1.0<br>[0.97 – 1.10] | 1* | 81.5 | 2 | -158 |
| no-Cas9_no-gRNAs | initial<br>off-target | viability | 162<br>[101 - 243] | 1* | 1.12<br>[0.84 – 1.63] | 81.5 | 2 | -158 |
| no-Cas9_no-gRNAs | neutral | none | 162<br>[101 – 243] | 1* | 1* | 81.5 | 1 | -161 |
| Cas9HF1_gRNAs | construct | viability | 396<br>[240 – 608] | 1.0<br>[0.97 – 1.04] | 1* | 88.1 | 2 | -171 |
| Cas9HF1_gRNAs | initial<br>off-target | viability | 433<br>[263 - 655] | 1* | 1.18<br>[0.99 – 1.45] | 89.7 | 2 | -175 |
| Cas9HF1_gRNAs | neutral | none | 396<br>[240 – 608] | 1* | 1* | 88.1 | 1 | -174 |

Each row shows the parameter estimates ( $\hat{N}_e$ = effective population size), maximum log Likelihood ( $\ln\hat{L}$ ), number of free parameters in the maximum likelihood framework ( $P$ ), and corrected Akaike Information Criterion value ( $AICc = 2p - 2\ln\hat{L} + (2p^2 + 2p)/(n - p - 1)$ where $n$ = number of generation transitions; $n= 20$ for Cas9\_no-gRNAs, no-Cas9\_no-gRNAs, and $n= 18$ for Cas9HF1\_gRNAs) for a specific construct, model and selection type. 1\* entries indicate that a parameter was fixed at 1 (= no fitness effect is estimated). Values in squared brackets in the parameter estimate columns represent the 95 % confidence intervals estimated from a likelihood ratio test with one degree of freedom.

### Phenotypic fitness assays

As a complementary validation of our fitness measurements from the cage experiments, we conducted three independent phenotypic assays (mate choice, fecundity, and viability) to estimate the fitness costs of the Cas9_gRNAs construct (see Supplementary Methods & Results). These assays broadly confirmed our previous findings. In particular, we found that Cas9_gRNAs homozygous males were 46.15 % less likely to be picked as mates by EGFP homozygous females (Figure S4A), and Cas9_gRNAs homozygous females laid on average 24.5% less eggs than EGFP homozygous females (Figure S4B). However, in contrast to our cage experiments where a model of viability-based fitness effect best matched the data (Table 1), we did not observe reduced viability for Cas9_gRNAs carrying flies in the individual assay (Figure S4C). However, this lack of difference in viability between EGFP and Cas9_gRNAs carrying flies could be explained by the assay environment. All phenotypic assays were conducted in vials, an environment where larvae experience much less resource competition than in the densely populated cage populations, which can significantly influence relative viability of different genotypes (44). Indeed, individuals that showed reduced fecundity or mating success in individual assays may have not survived to the adult stage in cage environments, representing a viability cost in that system. In addition, the viability assay examined only Cas9_gRNAs/EGFP heterozygotes, which may not have suffered from off-target effects to the full extent because they received "wild-type" off-target sites from one parent that did not carry the construct.

### Cas9HF1 homing drive

Our finding that off-target effects appear to be the primary driver for fitness costs from genomic Cas9 expression raises the question of whether Cas9HF1 would constitute a superior choice for gene drive strategies. As a proof-of-principle that Cas9HF1 is indeed a feasible alternative, we designed a homing drive that is identical to a previous drive (45), except that it uses Cas9HF1 instead of standard Cas9. This drive targets an artificial EGFP target locus with a single gRNA (see Methods).

We first crossed male flies carrying one of the two drives to females with the same EGFP target site used in our cage experiments. Individuals heterozygous for the homing drive and an EGFP allele were then further crossed to flies homozygous for EGFP, or to *w*^*1118*^ females for several of the male drive heterozygotes. The progeny of these crosses was phenotyped for DsRed, indicating presence of a drive allele, and EGFP, indicating the presence of an intact target allele (or more rarely, a resistance allele that preserved the function of EGFP). Disrupted EGFP alleles that did not show green fluorescence indicated the presence of a resistance allele.

We observed similar performance between the Cas9HF1 drive (Data Set S1) and the standard Cas9 drive (Data Set S2). Drive conversion efficiency for the Cas9HF1 drive was estimated at 80±2% for females and 59±3% for males, which was not significantly different than the rates for the standard Cas9 drive (83±2% for females and 61±2% for males) (*P* = 0.321 for female heterozygotes and *P* = 0.5513 for male heterozygotes, Fisher’s Exact Test). For both drives, all EGFP alleles in male heterozygotes that had not been converted to drive alleles were converted to resistance alleles, as indicated by the lack of EGFP phenotype in all progeny from crosses with *w^1118^* females. Both drives also had similar rates of resistance allele formation in the early embryo due to maternally deposited Cas9 (95±1% for alleles that disrupt EGFP for Cas9HF1, and 96±1% for standard Cas9, *P* = 0.3956, Fisher’s Exact Test). Together, these data demonstrate that homing drives with Cas9HF1 are capable of similar performance as drives using standard Cas9.

## Discussion

Negative selection will tend to displace any alleles from the population that are sufficiently deleterious. This effect can be quantified by an allele’s fitness, specifying the relative reproductive success between carriers and non-carriers of the allele. In this study, we measured the fitness of transgenic Cas9/gRNA alleles in *D. melanogaster*, which constitute an essential component of proposed applications such as CRISPR gene drives. A quantitative estimate of the fitness costs imposed by such constructs is critical for assessing the expected performance and limitations of these proposed applications.

Our constructs were designed to mimic a gene drive, yet without homing or any other mechanism that would facilitate super-Mendelian inheritance. This allowed us to estimate the “baseline” fitness costs of such systems. In particular, we inferred fitness by tracking allele frequencies in cage populations, which provides a powerful method for fitness estimation by integrating selective effects over all life stages and affected phenotypes (42). We did not observe detectable fitness costs due to Cas9/gRNA integration, expression, and on-target activity (which we refer to as “direct” costs) in our experiments. However, we did detect a moderate level of fitness cost resulting presumably from off-target effects. Such off-target costs were avoided when we used a high-fidelity version of Cas9 designed to minimize off-target cleavage (41). While the effects of off-target cutting in cells transiently exposed to Cas9 have already been extensively studied (46–49), our results demonstrate that such cleavage may have more substantial negative consequences over multiple generations when the cells are continuously expressing Cas9 in the germline from a genomic source.

Our study design does have some limitations that reduce the generality of our conclusions. First and foremost, off-target effects can vary substantially depending on the specific target sequence(s), genome composition, and expression patterns of Cas9 and gRNAs (15, 50). While we specifically selected gRNAs with a low number of predicted off-target sites to minimize such effects, this may not be possible for every application. Furthermore, the prediction of off-target sites may not always be accurate, potentially missing important sites. Some applications may also require the use of different promoters with higher somatic expression rates than *nanos* (22, 24, 28, 29), which could increase fitness costs caused by off-target cleavage. On the other hand, Cas9 expression may be lower in other organisms or at other genomic sites, and some applications might require fewer than the four gRNAs we included in our constructs, thereby potentially reducing off-target effects.

Another limitation is that our maximum likelihood framework for fitness estimation necessarily relied on a simplistic and highly idealized model. Most importantly, we modeled only a single genomic site to represent fitness costs from off-target cutting, and we used codominant fitness costs. In reality, there could be many off-target cut sites, with variable types of alleles after cleavage, different fitness costs, dominance relationships, degrees of genetic linkage, and possibly even epistatic interactions between them. Given the limited number of data points in our cage experiments, together with the large number of conceivable models, it is questionable whether our maximum likelihood framework could robustly infer the “correct” model.

The same holds true for the inference of different fitness components. While we did compare models where fitness affected viability versus models where it affected fecundity and mate choice, we believe that any conclusions from these comparisons should be taken with a grain of salt, given how many assumptions still went into each model (e.g., multiplicative fitness costs, same costs in males and females, and equal costs for fecundity and mate choice). Indeed, while our maximum likelihood analysis of the cage experiments ranked the viability model higher than the fecundity/mate choice model, we did observe reduced fecundity and mating success of genotypes carrying the Cas9_gRNAs construct in our phenotypic assays, while not finding a substantial effect on viability. However, as explained above, this could be due to a lower power of the phenotypic assays where only heterozygotes were studied, and/or the fact that many of those individuals with reduced fecundity and mating success in the phenotypic assays would not actually have survived into adult stage in the cage populations due to increased larval competition at higher densities.

Finally, we note that we did not attempt to identify the specific off-target cut sites that presumably caused the observed fitness effects and then track their allele frequencies. Such an analysis would have required time-resolved whole-genome sequencing on the population level. While potentially interesting, off-target sites would be construct-specific, and such an elaborate analysis may thus be more suitable when developing strategies to address off-target effects in a particular construct intended for release. Indeed, many studies have analyzed such individual off-target mutations in a large number of settings, including a study in mosquito gene drive (51). The focus of our study, however, was to elucidate the combined effects of such off-target cleavage on reproductive success, which has not been previously studied in the context of genomic integration of CRISPR elements. By directly measuring the “fitness” of a given construct on the population level, our approach is complementary to molecular studies that seek to identify all off-target mutations and then score their potential phenotypic effects. It also allows us to avoid the complexity of characterizing large numbers of potentially rare mutations, determining whether they are actually caused by off-target cleavage, and predicting what potential effect they may have on an organism’s fitness.

Though considerable uncertainty remains regarding the precise nature of the fitness effects of our constructs, the overall finding that genomic Cas9/gRNA-expression imposes a moderate fitness cost is robust. The fact that we only detected such costs for the construct that expressed both Cas9 and gRNAs, but not those lacking gRNAs, suggests that these costs are primarily due to cleavage activity, rather than just the expression of Cas9 or its genomic integration. Further, the fact that we did not observe any fitness costs when Cas9 was replaced with Cas9HF1 suggests that off-target effects are likely the driving factor for these costs. All of this is also consistent with the “bottoming out” phenomenon observed for construct frequencies in our cage experiments with Cas9_gRNAs, which can be explained by the accumulation of cut alleles at the off-target sites over the course of the experiment. It would be more difficult to reconcile with models where direct fitness costs are the driving factor or where fitness costs would be due to the specific genetic background of the construct flies or health-related effects of the initially released flies.

Our results have important implications for the modeling of gene drive approaches. Thus far, only direct fitness costs have been modeled in such studies, arising from the CRISPR nuclease itself, a payload gene, or cleavage of the intended target site. It is well known that such direct fitness costs can reduce the power of a suppression drive (38, 39, 52) and reduce the persistence of a modification drive in the face of resistance alleles (12). If such fitness costs are in fact lower than 5%, as suggested by our study in *Drosophila* at least, then they would not be expected to substantially impede the spread of suppression drives, though such drives could still suffer from other forms of direct fitness costs such as haploinsufficiency of the target gene and somatic Cas9 expression and cleavage. The direct fitness of modification drives would be largely determined by their cargo gene(s) and possibly their rescue efficiency if they involve use of a recoded gene (32, 40, 53, 54).

On the other hand, if off-target effects are in fact the primary driver of fitness costs of a drive with otherwise low direct fitness costs, as suggested by our study for *Drosophila* with the *nanos* promoter, this should result in different population dynamics. In a modification drive, such off-target effects would likely only slow the drive initially. After the drive has spread through most of the population, and cleaved off-target alleles had time to accumulate, resistance alleles would not be as selectively advantageous. Thus, it would take them much longer to outcompete drive alleles in the long run as compared to a scenario where direct fitness costs are the primary driver. A suppression drive may still suffer from cuts at off-target sites in a manner more closely resembling direct fitness costs, because these effects come into play during the early spread of a drive, often the most critical period in determining the fate of a suppression drive (38, 39, 52). However, if the rate at which off-target mutations form is sufficiently low, then mutated off-target sequences may not have a large effect on population dynamics. This has been shown in a recent study on a suppression drive with a germline-restricted promoter and a single gRNA that successfully eliminated a mosquito cage population before substantial amounts of off-target cleavage could occur (51).

We demonstrated that Cas9HF1, which largely eliminates off-target cleavage (41), does not induce substantial negative fitness effects when used as a replacement for standard Cas9 in our cage populations. Furthermore, we showed that homing drives with either form of Cas9 perform similarly. We therefore recommend that gene drives, as well as other applications that require the genomic integration of CRISPR endonucleases, should move from standard *Streptococcus pyrogenes* Cas9 to higher fidelity versions that can effectively minimize off-target effects. This would have the added advantage of reducing the generation of unanticipated genetic changes in natural populations from off-target cleavage and repair. One potential drawback of these nucleases is that they tend to have a lower cleavage rate that can depend on the specific gRNA sequence employed (55–60). In practice, this may reduce the number of available gRNA target sites and increase the need for initial evaluation of gRNA targets. However, newer improved forms of Cas9 (55, 56, 58–60), including ones with an expanded range of target sites (57), promise to ameliorate this issue.

In conclusion, we demonstrated that genomic CRISPR/Cas9 expression in *D. melanogaster* can impose a moderate level of fitness costs, most likely via off-target effects. Our results further indicate that fitness costs can be effectively minimized by using a high-fidelity endonuclease with reduced off-target cleavage. Future studies should investigate whether these conclusions hold in other experiments involving different constructs, target sites, and other organisms.

## Methods

### Plasmid construction

The starting plasmid pDsRed (Addgene plasmid #51019) was provided by Melissa Harrison, Kate O’Connor-Giles, and Jill Wildonger, pnos-Cas9-nos (61) (Addgene plasmid #62208) was provided by Simon Bullock, and VP12 (41) (Addgene plasmid #72247) was provided by Simon Bullock. Starting plasmids ATSacG, TTTgRNAtRNAi, TTTgRNAt, BHDgN1c, and BHDgN1cv3 were constructed in a previous study (45). Restriction enzymes for plasmid digestion, Q5 Hot Start DNA Polymerase for PCR, and Assembly Master Mix for Gibson assembly were acquired from New England Biolabs. Oligonucleotides and gBlocks were obtained from Integrated DNA Technologies. JM109 competent cells and ZymoPure Midiprep kit from Zymo Research were used to transform and purify plasmids. Cas9 gRNA target sequences were identified by the use of CRISPR Optimal Target Finder (62). A list of DNA fragments, plasmids, primers, and restriction enzymes used for cloning of each construct can be found in the Supplemental Information, together with annotated sequences of the final drive insertion plasmids (ApE format, http://biologylabs.utah.edu/jorgensen/wayned/ape).

### Generation of transgenic lines

Injections were conducted by Rainbow Transgenic Flies. The donor plasmid (Cas9_gRNAs, Cas9_no-gRNAs, no-Cas9_no-gRNAs, Cas9HF1_gRNAs, or BHDgNf1v2) (~500 ng/μL) was injected along with plasmid BHDgg1c (or TTTgU1 for BHDgNf1v2) (45) (~100 ng/μL), which provided additional gRNAs for transformation, and pBS-Hsp70-Cas9 (~500 ng/μL, from Melissa Harrison & Kate O’Connor-Giles & Jill Wildonger, Addgene plasmid #45945) providing Cas9. A 10 mM Tris-HCl, 100 μM EDTA solution at pH 8.5 was used for the injection. Most constructs were injected into *w*^*1118*^ flies, but BHDgNf1v2 was injected into flies with ATSacG (45). Transformants were identified by the presence of DsRed fluorescent protein in the eyes, which usually indicated successful construct insertion.

### Maintenance of transgenic flies with active Cas9HF1 gene drive

To minimize risk of accidental release, all flies with an active homing gene drive system were kept at the Sarkaria Arthropod Research Laboratory at Cornell University under Arthropod Containment Level 2 protocols in accordance with USDA APHIS standards. In addition, the synthetic target site drive system (30) prevents drive conversion in wild-type flies, which lack the EGFP target site. All safety standards were approved by the Cornell University Institutional Biosafety Committee.

### Experimental fly populations

The experimental fly populations were maintained on Bloomington Standard medium in 30×30×30 cm fly cages (Bugdorm). Flies were kept at constant temperature (25°C, 14 hours light, 10 hours dark), with non-overlapping generations. 0 – 2 day-old flies of one generation were allowed to lay eggs on fresh medium (8 food bottles per cage) for 24 hours. After that, the adults were frozen at −20°C for later phenotyping, and the new generation was allowed to develop for 11-12 days, before fresh medium was provided and a new generation cycle starts. The ancestral generation of each cage was generated by allowing homozygous EGFP flies and flies homozygous for the construct to deposit eggs for 24 hours separately from each other in four food bottles each. These eight egg-containing bottles were put in the fly cages to start one experimental fly population. Seven replicates of Cas9_gRNAs, and two replicates each for Cas9_no-gRNAs, no-Cas9_no-gRNAs, and Cas9HF1_gRNAs were maintained.

### Phenotyping experimental fly populations

The dominant fluorescent markers, EGFP and DsRed, allow a direct readout of the genotype by screening the fluorescent phenotype of an individual fly. Flies that are only red fluorescent are construct homozygotes, flies that are only green fluorescent do not carry any construct, and flies that are fluorescent for both colors carry one construct copy.

For each experimental population and generation, all individuals were screened for their genotypes using either a stereo dissecting microscope in combination with the NIGHTSEA system, or an automated image-based screening pipeline we specifically developed for this purpose. Quantifying phenotypic traits (e.g. pupae size, the amount of laid eggs) in an automated way has been done successfully before in *Drosophila* (63, 64). In our image-based screening pipeline, three pictures were taken for each batch of flies: a white light picture to determine the number and the position of the flies, one fluorescent picture filtered to screen for DsRed, and one fluorescent picture filtered to screen for EGFP expression.

We used a Canon EOS Rebel T6 with a 18-55 mm lens for image acquisition. The camera was held in a fixed position by a bracket 25 cm above the frozen flies spread on a black poster board. NIGHTSEA light heads (Green and Royalblue) were used as light sources. The light sources both for white and fluorescent light were covered with a paper tissue for diffusion. For the fluorescent pictures, barrier filters (Tiffen 58 mm Dark Red #29; Tiffen 58 mm Green #58) were used, attached with a magnetic XUME Lens/Filter system to the camera. Except for the filter change, the camera was fully controlled through a PC interface (EOS Utility 2 software). Focus was set automatically under white light and was kept constant for the fluorescent pictures. First, a white light picture (F 5.6, ISO 100, exposure time 1’’) was taken to determine the number and positions of the flies. Second, a picture under NIGHTSEA Green light with the Tiffen Dark red #29 filter (F 5.6, ISO 400, exposure time 30’’) was taken to determine, whether flies express DsRed. Third, a picture under NIGHTSEA Royal Blue with the Tiffen Dark Green #58 barrier filter (F 5.6, ISO 400, exposure time 25’’) was taken to screen flies for EGFP expression.

We used the ImageJ distribution Fiji (v 2.0.0-rc-69/1.52p) (65, 66) to process and analyze the picture sets with an in-house ImageJ macro: The three multi-channel images were split into the respective red, green, and blue image components. Further analysis included the red and the green image component of the white light picture, the red image component of the red fluorescent picture, and the green image component of the green fluorescent picture. The four remaining images were merged into a stack, and we performed slice alignment (matching method: normalized correlation coefficient) based on a selected landmark using the plugin Template_Matching.jar (67). We used a rectangular piece of white tape on the black poster board as landmark. To obtain the contours of the flies, we calculated the difference between the red and the green image component of the white light picture and applied a median and a Gaussian filter (radius = 3 pixels). After that, the picture was binarized using global thresholding (option: Max Entropy) (68). The binary image was post-processed (functions: Fill Holes, Open) before the position and the size of individual particles (=flies) were determined with the Analyze Particles method of ImageJ (minimum size = 750 pixels^2^). To account for translocations that have not been corrected for by the slice alignment (e.g., when the position of the fly changed slightly), the convex hull for each particle was calculated and enlarged by 20 pixels. A median filter (radius = 2 pixels) was applied to both fluorescent pictures before each particle (= fly) was scanned by a human investigator for the eye fluorescent pattern in both fluorescent pictures. We compared the image-based screening pipeline to the screening method using a stereo dissecting microscope and found that the estimated genotype frequencies deviate not more than 1% from each other (n = 646 flies, 4 picture sets).

### Phenotype data analysis, Cas9HF1 homing gene drive

When calculating drive parameters, we pooled offspring from the same type of cross together and calculated rates from the combined counts. A potential issue of this pooling approach is that batch effects could distort rate and error estimates (offspring were raised in separate vials with different parents). To account for such effects, we performed an alternate analysis as in previous studies (32, 45) by fitting a generalized linear mixed-effects model with a binomial distribution using the function glmer and a binomial link function (fit by maximum likelihood, Adaptive Gauss-Hermite Quadrature, nAGQ = 25). This allows for variance between batches, usually resulting in different rate estimates and increased error estimates. Offspring from a single vial were considered a distinct batch. This analysis was performed using the R statistical computing environment (v3.6.1) (69) with packages lme4 (1.1-21, https://cran.r-project.org/web/packages/lme4/index.html) and emmeans (1.4.2, https://cran.r-project.org/web/packages/emmeans/index.html). The R script we used for this analysis is available on Github (https://github.com/MesserLab/Binomial-Analysis). The results were similar to the pooled analysis and are provided in Supplementary Data Sets S1-S2.

### Genotyping

Flies were frozen, and DNA was extracted by grinding in 30 μL of 10 mM Tris-HCl pH 8, 1mM EDTA, 25 mM NaCl, and 200 μg/mL recombinant proteinase K (ThermoScientific), followed by incubation at 37°C for 30 minutes and then 95°C for 5 minutes. The DNA was used as a template for PCR using Q5 Hot Start DNA Polymerase from New England Biolabs. The region of interest containing gRNA target sites was amplified using DNA oligo primers AutoDLeft_S2_F and AutoDRight_S2_R. PCR products were purified after gel electrophoresis using a gel extraction kit (Zymo Research). Purified products were Sanger sequenced and analyzed with ApE (http://biologylabs.utah.edu/jorgensen/wayned/ape).

### Fitness cost estimation framework

To estimate the fitness costs of the different transgenic constructs in our *D. melanogaster* cage experiments, we modified a previously developed maximum likelihood inference framework (42). Specifically, we extended the original model to a two-locus model, where the first locus represents the construct insertion site and the second locus represents an idealized cut site. In this model, cleavage at the cut site could represent in principle the effects of non-specific DNA modifications (“off-target” effects) as well as the effects of cleavage at the desired gRNA target site (i.e., target site activity). However, the latter is not expected to impose any fitness costs for our constructs due to the intergenic location of the target site. Thus, we refer to the idealized cut site as “off-target” site. At the construct locus, the two possible allele states are EGFP/construct (observed by fluorescence); at the off-target site, the two possible states are uncut/cut (not directly observed). The two loci are assumed to be autosomal and unlinked. Thus, there are nine possible genotype combinations an individual could have in our model. Unless stated otherwise, we assumed that the construct homozygotes used for the ancestral generation of a cage are cut/cut homozygotes at the idealized off-target site. Since the construct is not homing, the allelic state of a single individual cannot change at the construct locus. By contrast, the allelic state at the off-target locus can be altered by cutting events in the germline or in the early embryo phase. Germline cutting will only impact the genotype of offspring in the next generation, while embryo cutting will directly change the individual’s genotype and hence expose it to any potential fitness effects of this new genotype. Both the germline and embryo off-target cut rates were set to 1 in our model. This means that any uncut allele at the off-target locus will be cut in the germline if the individual carries at least one construct allele (germline cute rate = 1). Furthermore, individuals will become cut/cut homozygotes if their mother carried a least one construct allele (embryo cute rate = 1; we assume that maternally deposited Cas9/gRNA is present in all such embryos).

A full inference model for the potential fitness costs of construct alleles and cut off-target alleles that includes all three previously implemented types of selection (mate choice, fecundity, viability) would feature a vast number of parameters that would be difficult to disentangle (42). For simplicity and to avoid overfitting, we therefore reduced model complexity with a series of assumptions: First, potential fitness costs were assumed to be equal for both sexes. Second, we either included only viability selection in the model, or included only mate choice (i.e., relative mating success for males with a particular genotype, reference value = 1) and fecundity selection (i.e., relative fecundity for females with a particular genotype, reference value = 1), both of equal magnitude. We further considered all fitness effects to be multiplicative across the two loci and for the two alleles at each locus (e.g., a construct homozygote would have a fitness equal to the square of a construct/EGFP heterozygote, given the same genotype at the off-target site). This results in two much more tractable inference models (viability and fecundity/mate choice) with only three parameters overall: the effective population size (*N*_*e*_), the relative fitness of construct/EGFP heterozygotes versus EGFP homozygotes (the “direct fitness parameter”), and the relative fitness of cut/uncut heterozygotes versus uncut homozygotes (the “off-target fitness parameter”).

## Supporting information

annotated sequences of final constructs

raw counts of each experimental population

picture sample set for the image-based screening pipeline

raw data of the phenotypic assays

## Availability of data and materials

The annotated sequences of the final construct insertions are available in ApE format (Supplemental file 1; constructs.zip). The raw counts of each experimental population (different constructs and the Cas9HF1 homing drive) can be found in Supplemental file 2 (Supplemental_Data_Sets.xlsx). The macro of the image-based screening pipeline is available on GitHub (https://github.com/MesserLab/CRISPR-Cas9-fitness-effects), a picture sample set for the image-based screening pipeline can be found in the Supplemental file 3 (example-pictures.zip). The raw data of the phenotypic assays can be found in Supplemental file 4 (phenotype_assays.zip). The maximum likelihood inference framework was implemented in R (v 3.6.0) (69), and is available together with all necessary scripts to reproduce the results on GitHub (https://github.com/MesserLab/CRISPR-Cas9-fitness-effects).

## Acknowledgements

This study was supported by the National Institutes of Health awards R21AI130635 to JC, AGC, and PWM, award F32AI138476 to JC, and award R01GM127418 to PWM. AML was supported by Vetmeduni Vienna Funds and an Austrian Science Funds grant (FWF; DK W1225-B20) awarded to Christian Schlötterer and an Austrian Marshall Plan Foundation fellowship. We thank Marlies Dolezal for helpful advice on the statistical analysis of the phenotypic assays. Special thanks to Charles Mazel from NIGHTSEA, and Guy Reeves for support developing the image-based screening pipeline.

## Supplemental Information

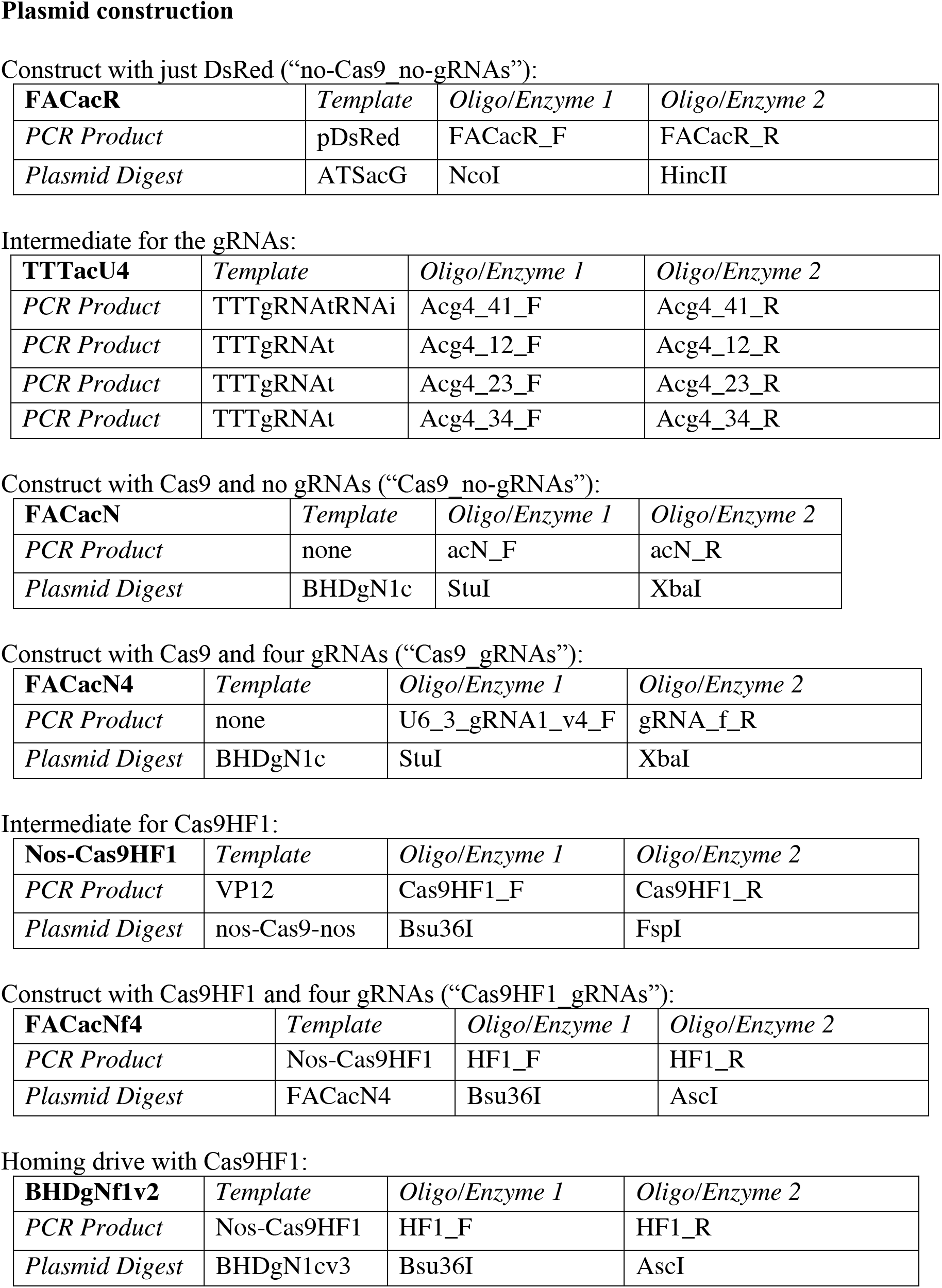
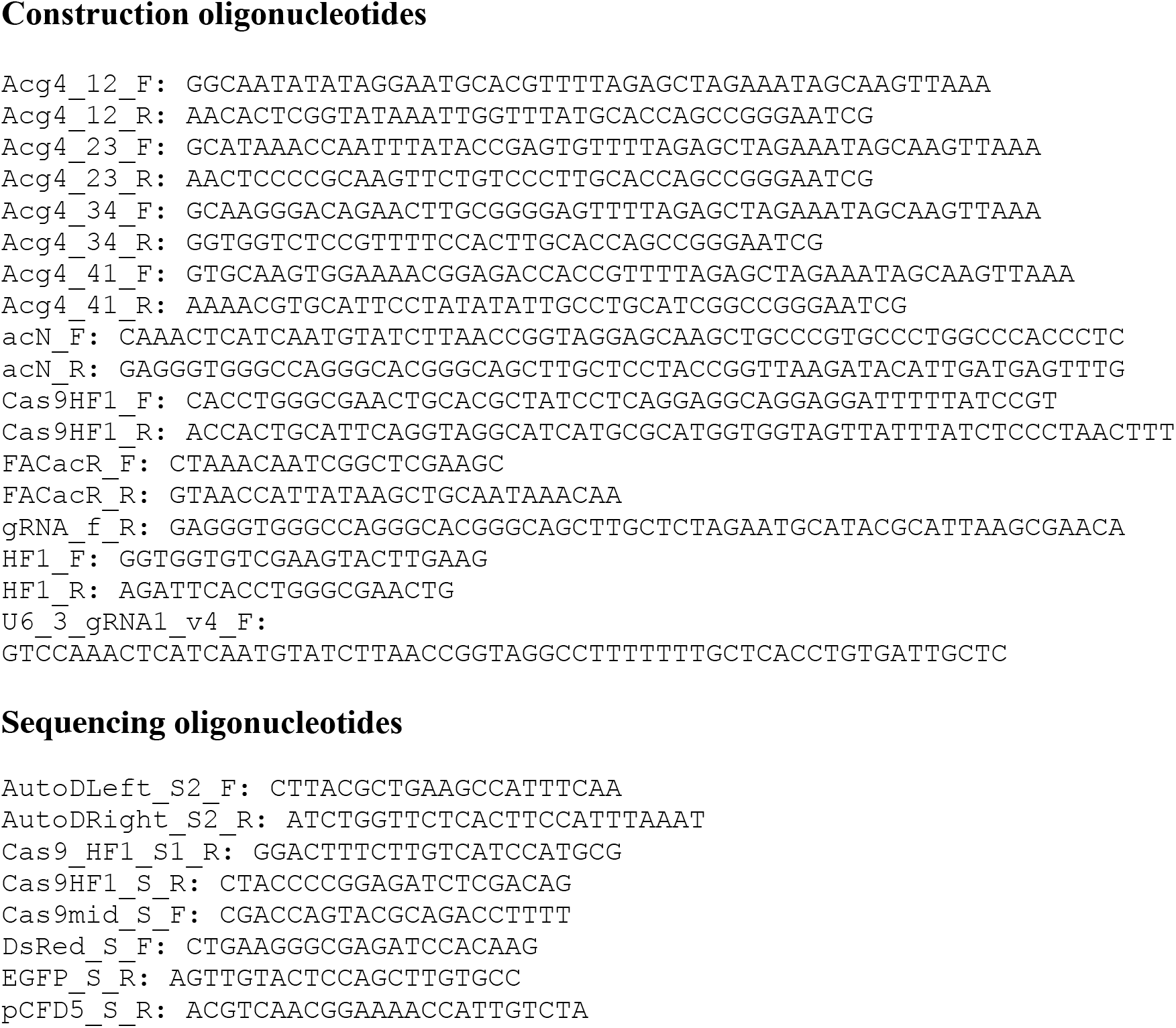

## Supplementary methods

### Phenotypic assays

We measured three fitness proxies for flies carrying Cas9_gRNAs constructs: mate choice, fecundity, and viability. All phenotypic assays were conducted on Bloomington Standard medium and under the same temperature (25°C) and light conditions (14 hours light/10 hours dark) as the caged populations. The statistical analysis of the phenotypic assays was conducted in R (v3.6.0). (69)

#### Mate choice

We conducted a mate choice assay to test for mating preferences of EGFP homozygous females. Individual 2-day-old virgin EGFP homozygous females were set up with one EGFP homozygous male and one Cas9_gRNAs homozygous male of the same age in a vial. After 24 hours, the adult flies were removed, and the genotypes of the eclosed offspring were assessed after 11 to 12 days. If the EGFP homozygous female has mated only with the male of the same genotype, only homozygous offspring is expected. We tested for deviations from an expected equal frequency of offspring genotypes under the null hypothesis of no mate preference via a binomial test.

#### Fecundity

We assessed the fecundity of EGFP homozygous, heterozygous, and Cas9_gRNAs homozygous females in individual crosses with EGFP homozygous males. Each individual single 2-day-old virgin female of a distinct genotype was crossed with one EGFP homozygous male of the same age. Crosses were flipped on fresh medium every 24 hours, and eggs were counted manually using a stereo dissecting microscope. Fecundity was defined as the total number of laid eggs per female over three consecutive days. To assess the impact of female genotype we fitted a linear model using function lm() with fecundity as response. The female genotype was the only fixed effect in the model. The residuals were both normally distributed and showed variance homogeneity, meeting all assumptions of a linear model. None of the used model diagnostics (Cook’s distance, DFbetas, leverage (70); calculated with the R package car (v3.0-3) (71)) indicated strongly influential cases or outliers. We used the R package emmeans (v1.4.7) (72) to conduct pairwise comparisons of the three assessed female genotypes.

#### Viability

We measured relative viability of heterozygous offspring of single crosses between EGFP homozygous males and heterozygous females. Single 2-day-old heterozygous virgin females were crossed each with one EGFP homozygous male of the same age. After 24 hours, the adult flies were removed, and the genotypes of the eclosed offspring was assessed after 11 to 12 days. Relative heterozygote viability was defined as the fraction of heterozygous offspring out of the total number of offspring, ranging between 0 and 1. If the genotype does not influence viability, we expect a relative heterozygote viability of 0.5. Relative heterozygote viability was tested for normality with an Anderson-Darling test (function ad.test() in the R package nortest (v1.0-4) (73)). We then used a one-sample t-test against a population mean of 0.5 for heterozygotes viability.

## Supplementary results

### Population cage experiments setups

All experiments started by crossing construct and EGFP homozygotes, except for replicates 1, and 2 of Cas9_gRNAs. These two experimental populations were set up with all three genotypes that originated from the same batch that included heterozygotes and both homozygotes. While construct homozygotes of Cas9_no-gRNAs, no-Cas9_no-gRNAs, and Cas9HF1_gRNAs were of the same age as the EGFP homozygotes they were mixed with to start the experiments, the age differed between EGFP and construct homozygotes for Cas9_gRNAs replicates 1, 2, 5, 6, and 7. To avoid confounding maternal effects on the construct frequency dynamics, we excluded for each of these replicates the first generation from the analysis. The full data set including the removed time points can be found in Supplemental File 2.

Population sizes were controlled via the limited egg-lay time period, which led to fluctuations in the number of flies per generation (Figure S1). Some experiments experienced bottlenecks due to high variation in food moisture content (resulting in either high or low larvae density).

### Phenotypic assays

#### Mate choice

We assessed the mate choice of 40 independent EGFP homozygous females that were each set up with one EGFP homozygous and one Cas9_gRNAS homozygous male in one single vial. 38 samples had exclusively EGFP homozygous or heterozygous offspring, whereas 2 samples displayed offspring of both genotypes. As this suggests multiple matings of the female, we excluded these two data points from the analysis. The estimated frequency of 0.684 of EGFP homozygous females choosing EGFP homozygous males (*n*=26) over Cas9_gRNAs homozygous males (*n*=12) as mates was significantly different from 0.5 (exact binomial test *P*=0.033; Figure S4A).

#### Fecundity

In total, we measured the fecundity of 128 independent females (27 EGFP homozygotes, 55 heterozygotes, and 46 Cas9_gRNAs homozygotes). Overall, the female genotype significantly influenced fecundity, which was defined as the total number of laid eggs per female over the course of three consecutive days (full-null model comparison *F*_2,125_=5.885, *P*=0.004). Cas9_gRNAs homozygous females are significantly less fecund than EGFP homozygous females. However, no significant difference was detected between EGFP homozygotes and heterozygotes, or heterozygotes and Cas9_gRNAs homozygotes respectively (Figure S4B).

#### Viability

We determined the relative heterozygote viability (=fraction of heterozygous offspring out of the total number of offspring, ranging between 0 and 1) of 56 independent fly crosses. We observed that the relative heterozygote viability is normally distributed (mean = 0.486, standard deviation = 0.098; *A*=0.405, *P*=0.343) and average heterozygote viability does not significantly differ from 0.5 (*t*_55_=−1.057, *P*=0.295; Figure S4C).

---

**Table S1.**
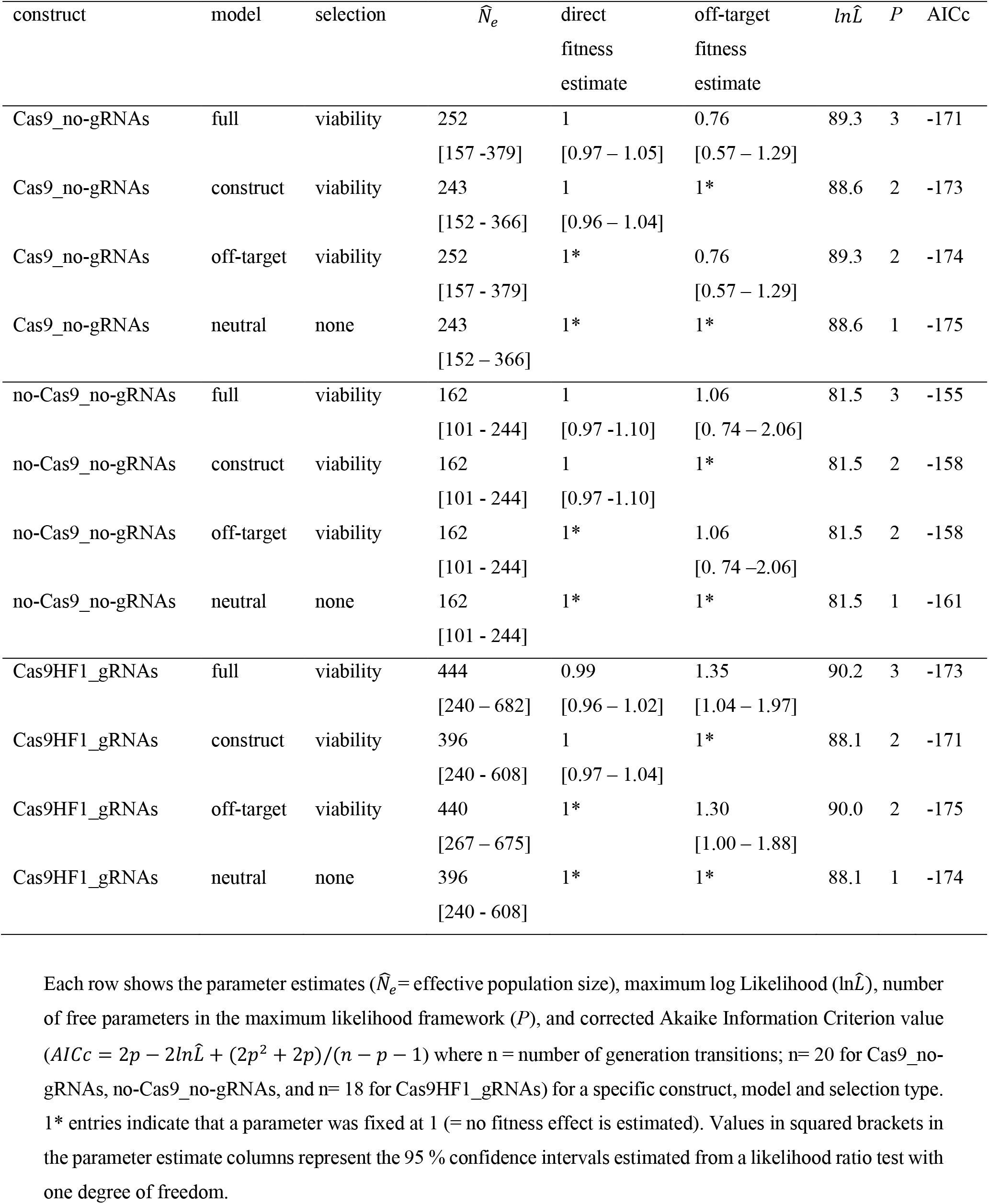
Model comparison of Cas9_no-gRNAs, no-Cas9_no-gRNAs, Cas9HF1_gRNAs – all cut parameters set to 1

| construct | model | selection | $\hat{N}_e$ | direct<br>fitness<br>estimate | off-target<br>fitness<br>estimate | $\ln\hat{L}$ | $P$ | AICc |
| --- | --- | --- | --- | --- | --- | --- | --- | --- |
| Cas9_no-gRNAs | full | viability | 252<br>[157 - 379] | 1<br>[0.97 – 1.05] | 0.76<br>[0.57 – 1.29] | 89.3 | 3 | -171 |
| Cas9_no-gRNAs | construct | viability | 243<br>[152 - 366] | 1<br>[0.96 – 1.04] | 1* | 88.6 | 2 | -173 |
| Cas9_no-gRNAs | off-target | viability | 252<br>[157 - 379] | 1* | 0.76<br>[0.57 – 1.29] | 89.3 | 2 | -174 |
| Cas9_no-gRNAs | neutral | none | 243<br>[152 – 366] | 1* | 1* | 88.6 | 1 | -175 |
| no-Cas9_no-gRNAs | full | viability | 162<br>[101 - 244] | 1<br>[0.97 -1.10] | 1.06<br>[0. 74 – 2.06] | 81.5 | 3 | -155 |
| no-Cas9_no-gRNAs | construct | viability | 162<br>[101 - 244] | 1<br>[0.97 -1.10] | 1* | 81.5 | 2 | -158 |
| no-Cas9_no-gRNAs | off-target | viability | 162<br>[101 - 244] | 1* | 1.06<br>[0. 74 –2.06] | 81.5 | 2 | -158 |
| no-Cas9_no-gRNAs | neutral | none | 162<br>[101 - 244] | 1* | 1* | 81.5 | 1 | -161 |
| Cas9HF1_gRNAs | full | viability | 444<br>[240 – 682] | 0.99<br>[0.96 – 1.02] | 1.35<br>[1.04 – 1.97] | 90.2 | 3 | -173 |
| Cas9HF1_gRNAs | construct | viability | 396<br>[240 - 608] | 1<br>[0.97 – 1.04] | 1* | 88.1 | 2 | -171 |
| Cas9HF1_gRNAs | off-target | viability | 440<br>[267 – 675] | 1* | 1.30<br>[1.00 – 1.88] | 90.0 | 2 | -175 |
| Cas9HF1_gRNAs | neutral | none | 396<br>[240 - 608] | 1* | 1* | 88.1 | 1 | -174 |

Each row shows the parameter estimates ( $\hat{N}_e$ = effective population size), maximum log Likelihood ( $\ln\hat{L}$ ), number of free parameters in the maximum likelihood framework ( $P$ ), and corrected Akaike Information Criterion value ( $AICc = 2p - 2\ln\hat{L} + (2p^2 + 2p)/(n - p - 1)$ where $n$ = number of generation transitions; $n= 20$ for Cas9\_no-gRNAs, no-Cas9\_no-gRNAs, and $n= 18$ for Cas9HF1\_gRNAs) for a specific construct, model and selection type. 1\* entries indicate that a parameter was fixed at 1 (= no fitness effect is estimated). Values in squared brackets in the parameter estimate columns represent the 95 % confidence intervals estimated from a likelihood ratio test with one degree of freedom.

**Figure S1.**
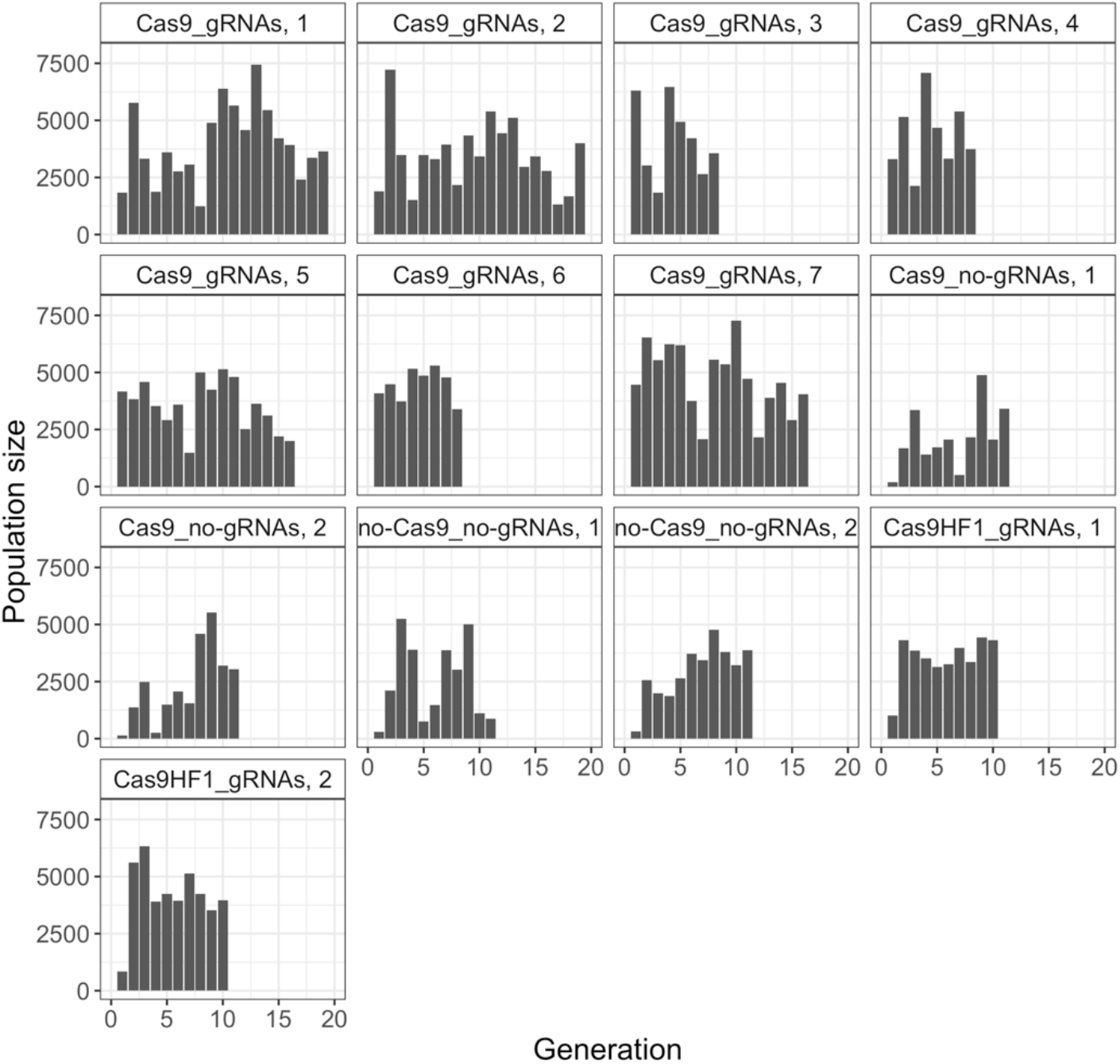
Population sizes of all *D. melanogaster* cage populations.

**Figure S2.**
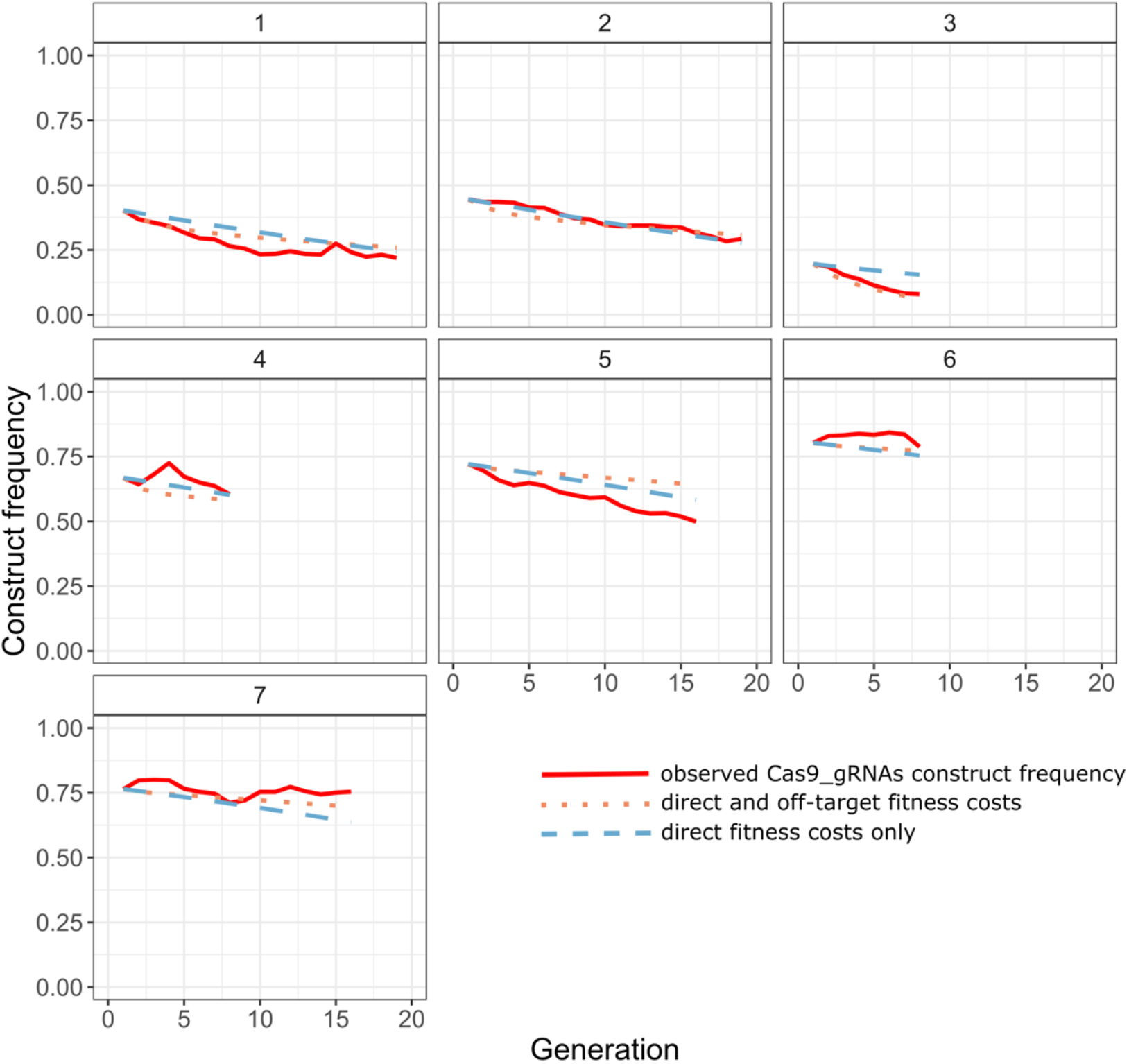
Comparison of observed construct frequencies (solid red line) in our experimental Cas9_gRNAs cages with the predicted trajectories of the full inference model with viability selection (dotted, orange line; off-target fitness = 0.84, direct fitness = 0.98), and the construct model with viability selection (dashed, blue line; direct fitness = 0.96), using the inferred maximum likelihood parameter estimates (Table 1). Genetic drift was not simulated.

**Figure S3.**
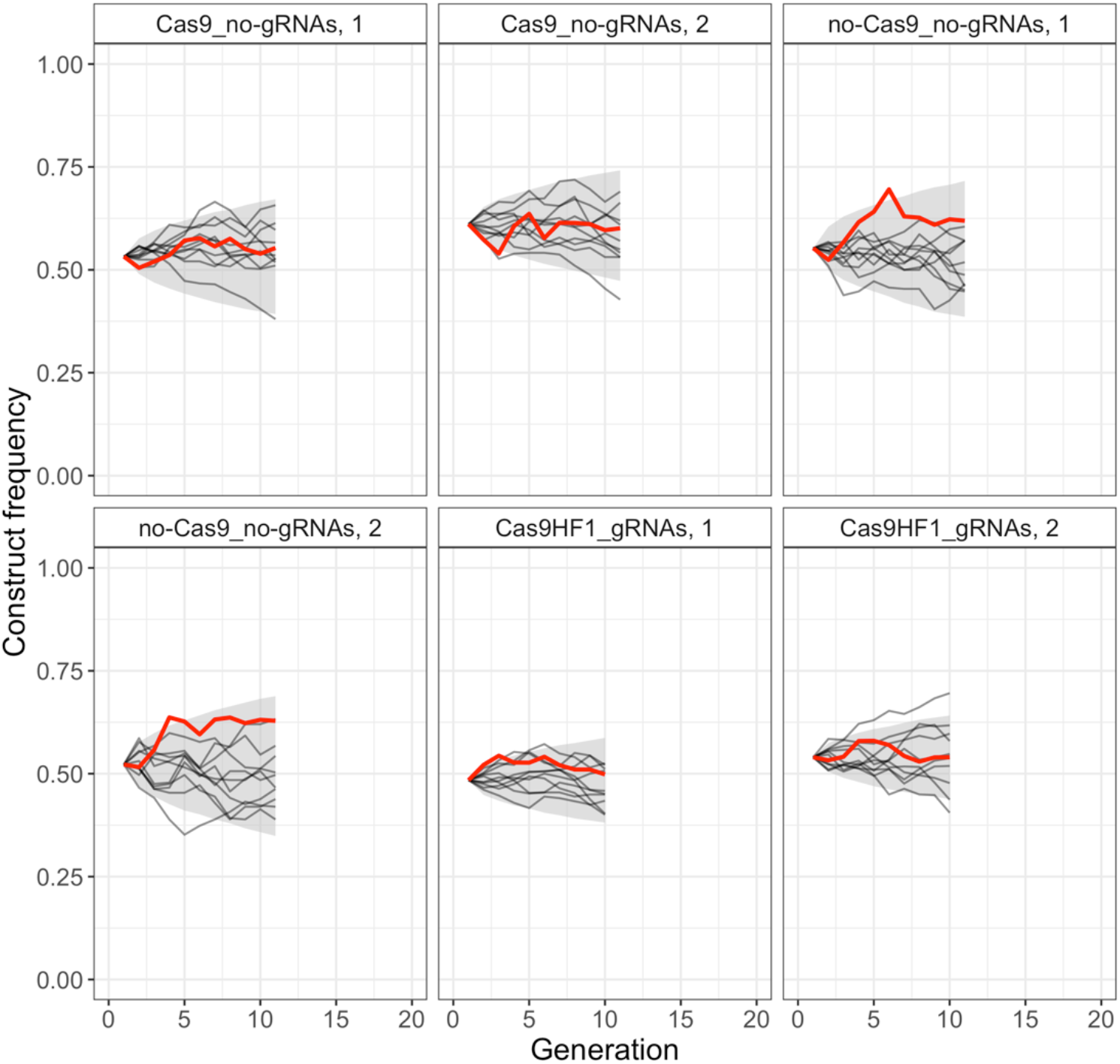
Comparison of observed construct frequencies with simulated trajectories of a neutral model 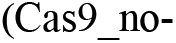 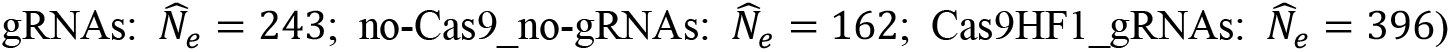. Solid red lines present observed construct frequencies, black lines show ten simulated trajectories for each cage, and the shaded area represents the range between the 2.5 and 97.5 percentile of the simulated trajectories (10,000 simulations per cage).

**Figure S4.**
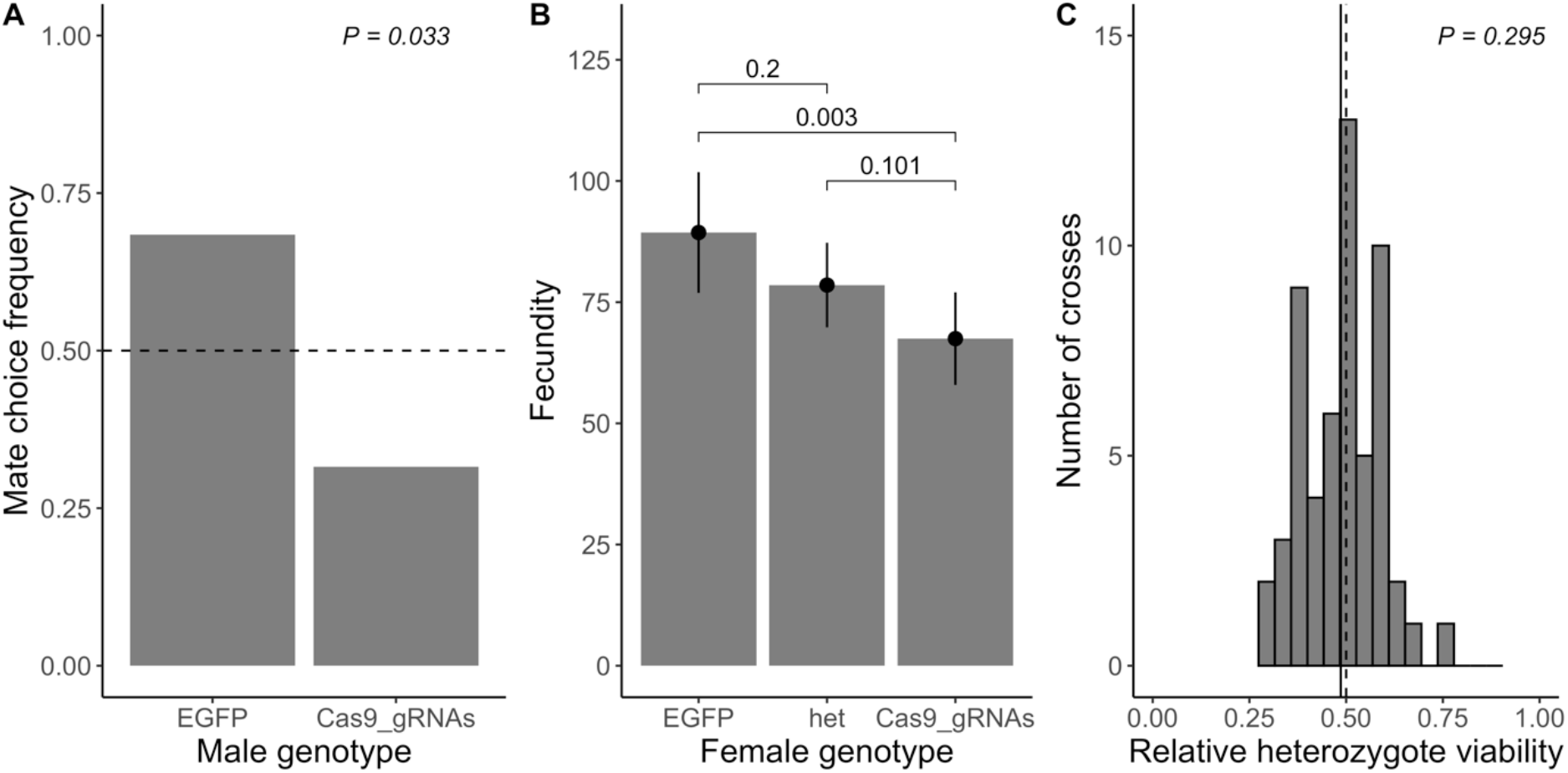
Direct measurement of fitness parameters. (A) Observed mate choice frequency (y-axis) of EGFP homozygous females choosing between EGFP and Cas9_gRNAs homozygous males (x-axis; as only two genotypes were tested, the frequencies sum up to 1). In case of no mate choice preference, the expected mate choice frequency is 0.5 (horizontal dashed line). The observed mate choice frequency of EGFP homozygous males was significantly different from 0.5 (exact binomial test; *P* = 0.033). (B) Average fecundity (y-axis) for each female genotype (x-axis). The observed average fecundity (= total number of eggs per female laid over the course of three consecutive days) is plotted for each female genotype separately (EGFP = EGFP homozygous females, het = heterozygous females, Cas9_gRNAS = Cas9_gRNAs homozygous females). All females were mated in individual crosses to EGFP homozygous males of the same age. The fitted model is shown as black dots with error bars displaying the 95 % confidence interval. P-values of pairwise genotype comparisons adjusted with the Tukey method are displayed above the bars. (C) Relative heterozygote viability (= fraction of heterozygous offspring of crosses between heterozygous females and EGFP homozygous males). If the genotype does not influence viability, we expect a relative heterozygote viability of 0.5 (vertical dashed line). The observed average heterozygote viability (vertical solid line) does not differ from 0.5 (one sample t-test; *P* = 0.295).

