## Supplementary material for "Fitness effects of CRISPR endonucleases in *Drosophila melanogaster* populations": annotated sequences of final constructs: README.txt

AutosomeC.ape	insertion site for EGFP on chromosome 2L
AutosomeD.ape	construct target site on chromosome 3L
BHDgNF1v2.ape	Cas9HF1 homing drive 
FACacN.ape		Cas9_no-gRNAs construct
FACacN4.ape		Cas9_gRNAs construct
FACacNf4.ape	Cas9HF1_gRNAs construct
FACacR.ape		no-Cas9_no-gRNAs construct
