## Supplementary material for "Fitness effects of CRISPR endonucleases in *Drosophila melanogaster* populations": picture sample set for the image-based screening pipeline: README.txt

IMG_001.JPG white light picture 
IMG_002.JPG fluorescent picture to check for DsRed
IMG_003.JPG fluorescent picture to check for EGFP 

Flies are grouped by genotype

Top row: eGFP homozygotes (n=24)
Middle row: heterozygots (n=90)
Bottom row: dsRed homozygotes (n=40)

For Macro usage, load the pictures in the following order:
1.) IMG_001.JPG
2.) IMG_002.JPG
3.) IMG_003.JPG

Macro requires Template Matching Plugin
https://sites.google.com/site/qingzongtseng/template-matching-ij-plugin#downloads
