## Supplementary figures and images for "Fitness effects of CRISPR endonucleases in *Drosophila melanogaster* populations"

### IMG_0001.JPG

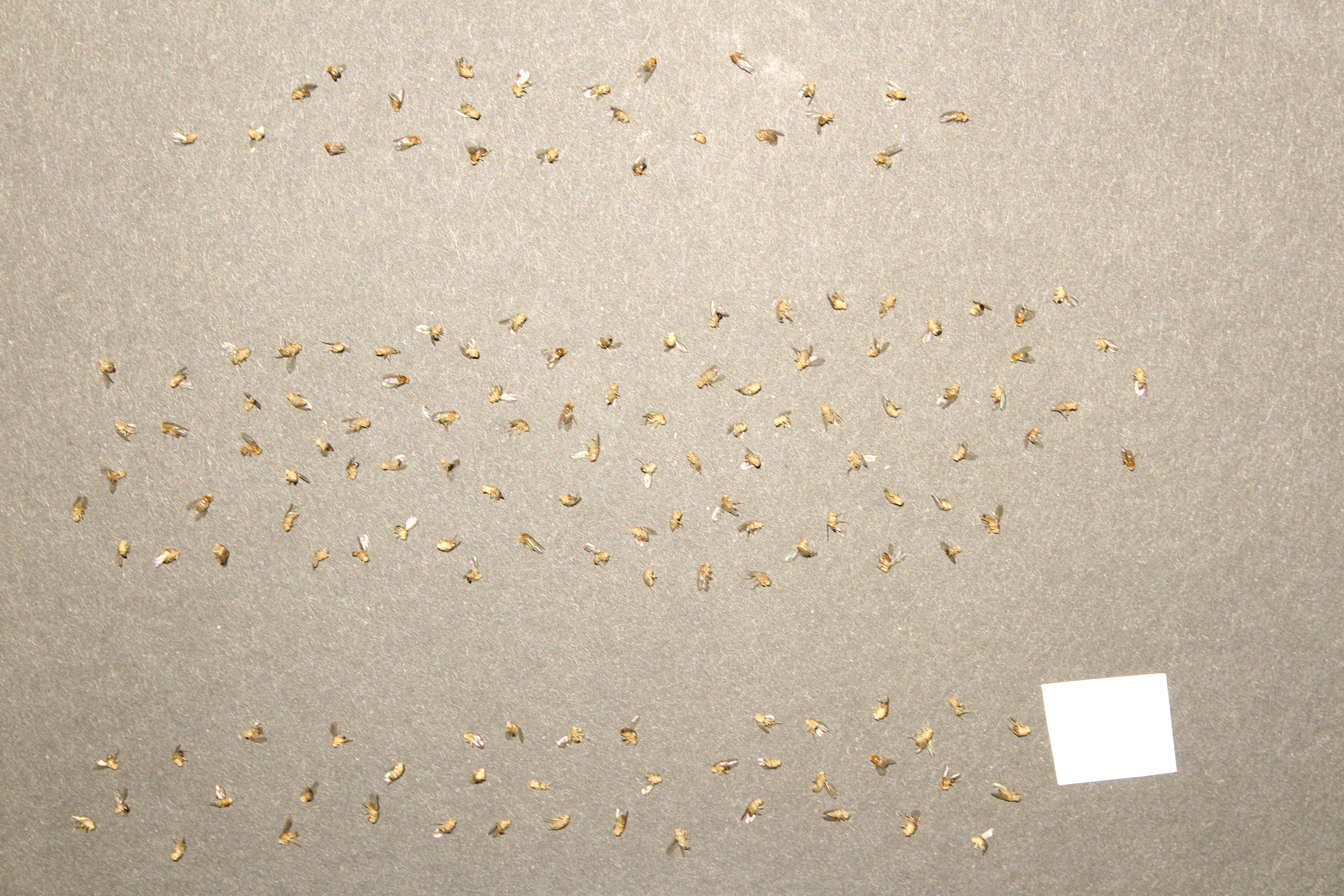

### IMG_0002.JPG

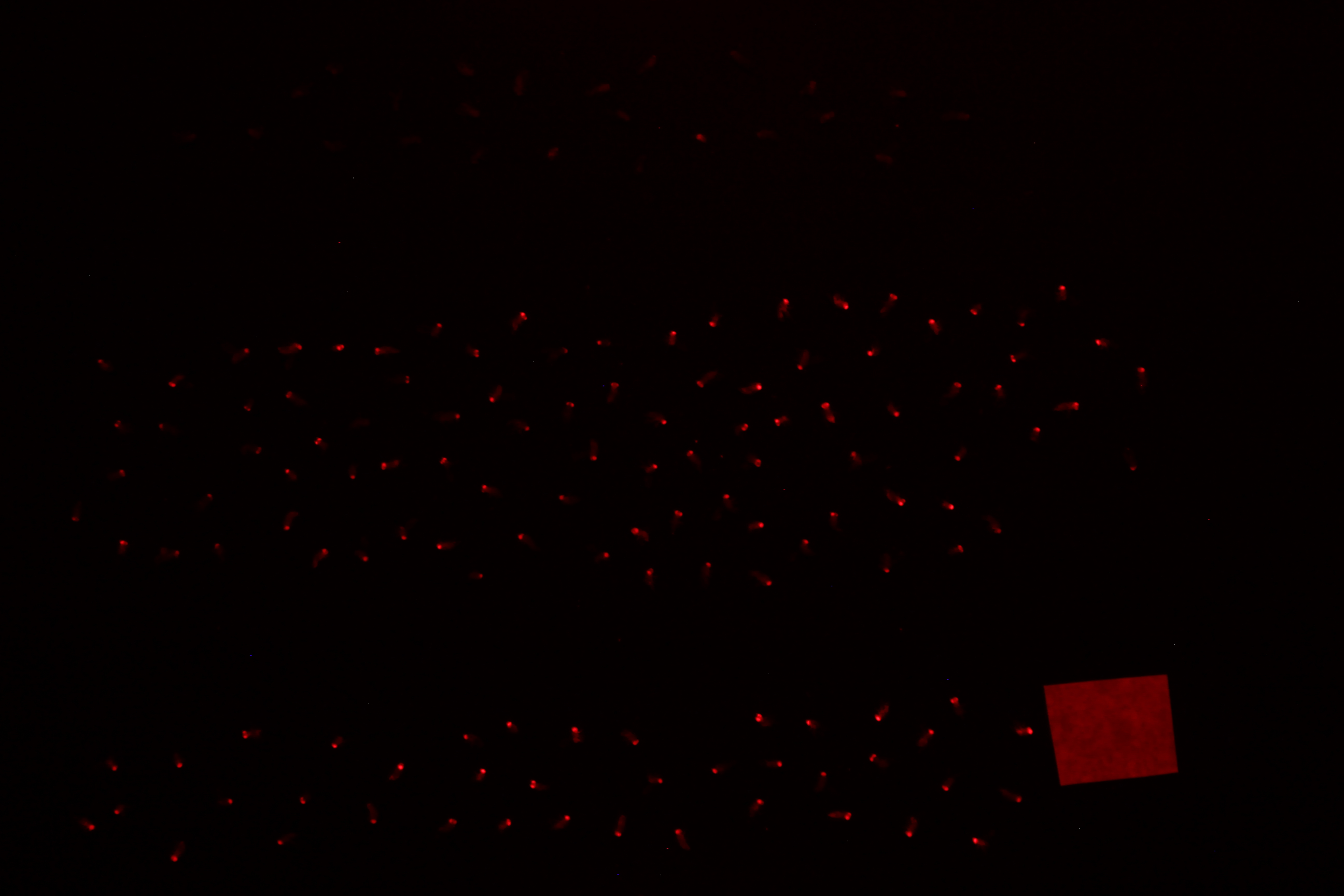

### IMG_0003.JPG

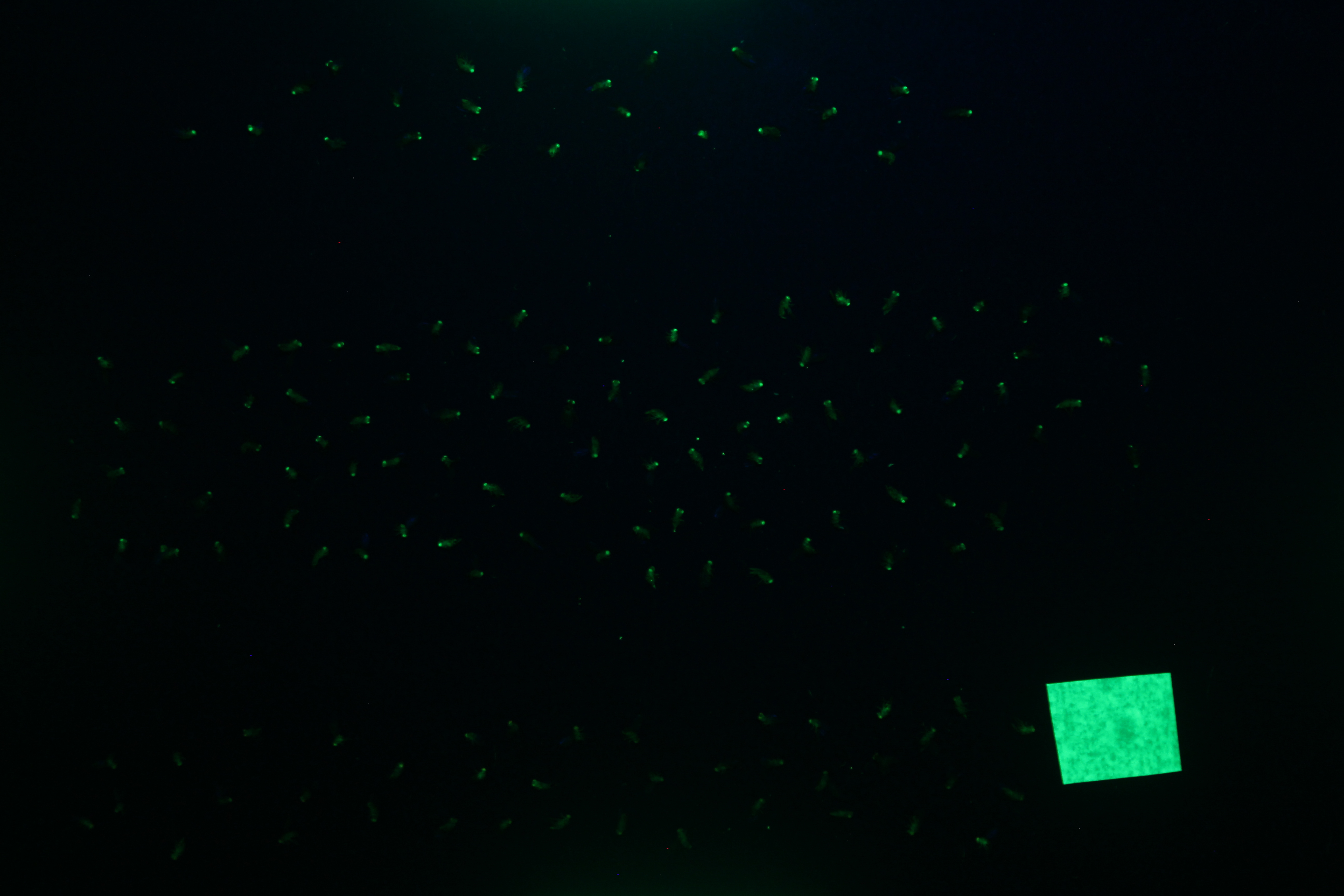
